# Tolerogenic Dendritic Cells Shape a Transmissible Gut Microbiota that Protects from Metabolic Diseases

**DOI:** 10.1101/2020.10.22.350033

**Authors:** Emelyne Lécuyer, Tiphaine Le Roy, Aurélie Gestin, Amélie Lacombe, Catherine Philippe, Maharajah Ponnaiah, Jean-Baptiste Huré, Magali Fradet, Farid Ichou, Samira Boudebbouze, Thierry Huby, Emmanuel Gautier, Moez Rhimi, Emmanuelle Maguin, Nathalie Kapel, Philippe Gérard, Nicolas Venteclef, Michèle Garlatti, Benoit Chassaing, Philippe Lesnik

**Author notes:** Corresponding authors: Dr Philippe Lesnik,. Dr Emelyne Lécuyer.

## Abstract

Excess of chronic contact between microbial motifs and intestinal immune cells are known to trigger a low-grade inflammation involved in many pathologies such as obesity and diabetes.

The important skewing of intestinal adaptive immunity in the context of diet-induced obesity (DIO) is well described but how dendritic cells (DCs) participate to these changes is still poorly documented. To address this question, transgenic mice with enhanced DCs lifespan and immunogenicity (DC^hBcl-2^ mice), are challenged with a high fat diet.

Those mice display resistance to DIO and metabolic alterations. The DIO resistant phenotype is associated with healthier parameters of intestinal barrier function and lower intestinal inflammation. DChBcl^-2^ DIO-resistant mice demonstrate a particular increase in tolerogenic DC numbers and function which is associated with strong intestinal IgA, Th17 and T regulatory immune responses.

Microbiota composition and function analyses reveal that the DC^hBcl-2^ mice microbiota is characterized by a lower immunogenicity and an enhanced butyrate production. Cohousing experiments and fecal microbial transplantations are sufficient to transfer the DIO resistance status to WT mice demonstrating that maintenance of DCs tolerogenic ability sustains a microbiota able to drive DIO resistance. DCs tolerogenic function is revealed as a new potent target in metabolic diseases management.

## Introduction

Metabolic syndrome consists in a pathophysiological state combining several metabolic abnormalities such as abdominal obesity, insulin resistance, hyperglycemia, dyslipidemia, hypertension and fatty liver (O’Neill and O’Driscoll 2015). The World Health Organization estimates that around 34% of the population has developed or is at risk to develop this syndrome that predisposes to cardiovascular diseases and cancers (Saklayen 2018).

A major component that triggers the initiation and the worsening of metabolic syndrome is chronic low-grade inflammation. Loss of homeostatic intestinal immunity leading to impaired intestinal barrier function has been described as a first step that precedes and predisposes to systemic low-grade inflammation associated with obesity and related metabolic disorders (Ding et al. 2010; Luck et al. 2015)(Luck et al. 2015). Modifications in the composition of the intestinal microbiota induced by dietary changes have been shown to trigger the development and the maintenance of many pathologies including metabolic disorders (Ding et al. 2010; Luck et al. 2015; Garidou et al. 2015; Winer et al. 2016; Petersen et al. 2019). The constant dialog between the microbiota and the immune system, essential for intestinal immune development at birth, is also critical in regulating the structure and composition of the intestinal microbiota throughout life (Brown, Sadarangani, and Finlay 2013).

In the last decades, many progresses have been made to better characterize the role of pro-inflammatory immune responses in the pathogenesis of metabolic dysfunctions (Winer et al. 2016; Fernández-Ruiz 2016). Among immune cells, few studies have focused on the role of intestinal antigen presenting cells (APCs) (Kawano et al. 2016; Zlotnikov-Klionsky et al. 2015). APCs such as dendritic cells (DCs) act as sensors of their environment, allowing them to be highly responsive to catch and transduce extracellular signals as antigens and cytokines. Stimulated-DCs can migrate into the draining lymph nodes where they prime the adaptive immune cells promoting either tolerance or pro-inflammatory immune responses (Coombes and Powrie 2008). The orientation between pro-inflammatory or tolerogenic adaptive immune priming is based on the signals they received and also depends on the maturation/activation status of the DCs. Detection of enteric pathogens through pattern recognition receptors (PRRs) induces DCs maturation, triggering pro-inflammatory adaptive immunity for pathogens clearance. Conversely, at steady state, mucosal DCs are known to promote tolerogenic immunity (Sun et al. 2007; Pabst and Mowat 2012). The retinaldehyde dehydrogenases (RALDHs) are key enzymatic activities related to DCs tolerogenic function and intestinal CD103^+^ RALDH^+^ DCs are described as mainly involved in this process (Mora et al. 2006; Chang, Ko, and Kweon 2014). After migration from the intestine into the draining lymph nodes, RALDH activity in DCs promotes IgA class-switching plasma cells, IL-17-producing T CD4^+^ (Th17) and IL-10-producing Foxp3^+^ T CD4^+^ (Treg). Moreover, tolerogenic DCs promote gut tropism marker on primed adaptive immune cells enabling them to migrate to the intestinal compartment to locally establish their tolerogenic functions. Several environmental cues are responsible for the maintenance of DCs tolerogenic function. Particular attention has been paid to decipher how microbiota-derived metabolites may influence tolerogenic activity of DCs. Butyrate is one short chain fatty acid (SCFA) described to orientate tolerogenic activity in mucosal DCs (Kaisar et al. 2017; Qiang et al. 2017).

The constant dialog between the microbiota and the immune system, essential for intestinal immune development at birth, is also critical in regulating the structure and composition of the intestinal microbiota throughout life (Brown, Sadarangani, and Finlay 2013). However, the contribution of DCs-microbiota crosstalk in orchestrating the progression of low-grade inflammation is unclear.

To decipher the impact of DCs, we used a mouse model developed in our laboratory for which the human anti-apoptotic factor B-cell lymphoma 2 (Bcl-2) has been inserted under the CD11c promoter to target DCs (DC^hBcl-2^ mice). Targeting Bcl-2 in DCs is known to promote increased DCs lifespan and this is associated with a higher DCs number in lymphoid organs. This strategy boosts adaptive immune responses in vitro (Nopora and Brocker 2002). In vivo, upon acute LPS challenge, the survival of DC^hBcl-2^ mice is enhanced and associated with increased DCs survival and increased capacity of DCs to prime adaptative immune responses related to Th1, Th17 and Tr1 polarization (Gautier et al. 2008; Gautier Emmanuel L. et al. 2009). Since it has been hypothesised that inappropriate intestinal Th polarization could promote the deleterious pro-inflammatory immune environment associated with metabolic alterations (Winer et al. 2016), we assessed how the increase DCs lifespan impacts on DC-microbiota crosstalk and host metabolism in the context of diet-induced obesity.

## Results

### DC^hBcl-2^ mice demonstrate resistance to HFD induced obesity and associated metabolic alterations

To decipher how DCs orchestrate diet-mediated obesity and metabolic alterations we used a mouse model enriched for DCs. This mouse model overexpresses the antiapoptotic human gene *hBcl2* under the control of a DC-related promoter CD11c. Despite the CD11c promoter is non-restrictive to DC, the hBcl2 transgene has been shown to be particularly expressed in CD11c^+^ DCs (Gautier et al. 2008). These DC^hBcl-2^ mice are characterized by enhanced DCs lifespan leading to increased immunogenicity affecting both pro-inflammatory and regulatory adaptive immunity (Th1, Th17 and Tr1 cells) (Gautier et al. 2008; Gautier Emmanuel L. et al. 2009).

We challenged adult WT and DC^hBcl-2^ mice with 60% fat diet (HFD) or control chow diet (CCD) upon 24 weeks. Although there was no difference in weight between WT and DC^hBcl-2^ mice on CCD (data not shown), DC^hBcl-2^ mice gained 10% less weight than WT mice after 12 weeks of HFD (Figure 1A). Those differences in weight gain were maintained until 24 weeks of HFD (Figure 1A,B).

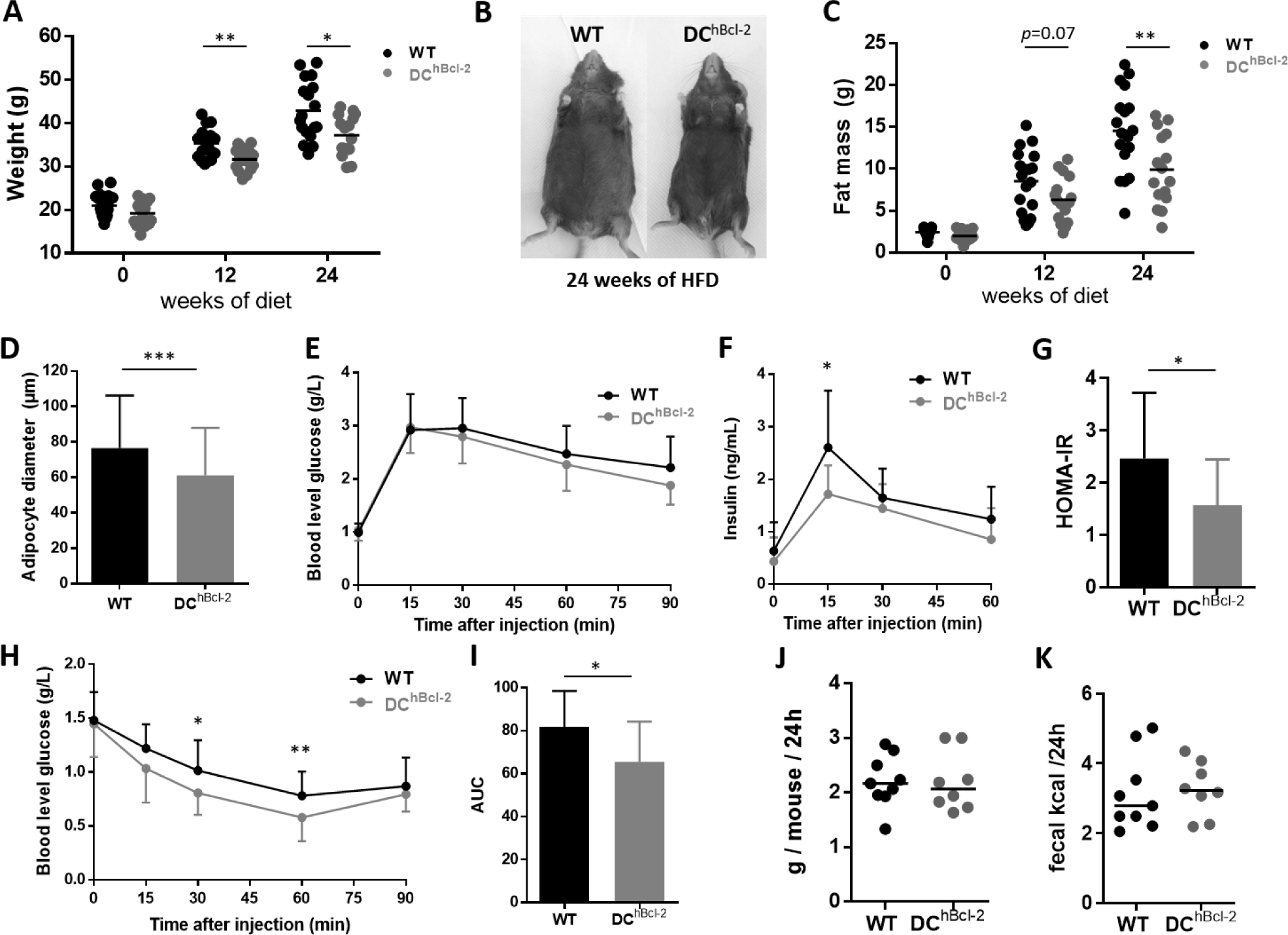

We wondered if changes in body weight corresponded with changes in adiposity. Body mass composition analysis all along the HFD indicated that despite the same increase in lean mass (figure supplement 1A), DC^hBcl-2^ mice gained less fat mass than their WT counterparts reaching at the end of the challenge 9.9 ± 4.2 g of fat compared to 14.5 ± 4.9 g as respective mean fat mass ± standard deviation (SD) (Figure 1C). The lower fat mass gain in DC^hBcl-2^ mice was associated with reduced adipocyte size (Figure 1D).

Regarding the differences in body fat composition we investigated how glucose metabolism was impacted in both WT and DC^hBcl-2^mice. After 13 weeks of HFD we performed an oral glucose tolerance test (OGTT). Despite comparable blood glucose levels (Figure 1E), we observed significant differences of circulating insulin levels following glucose administration (Figure 1F). Homeostasis Model Assessment of Insulin Resistance (HOMA-IR) indicated that DC^hBcl-2^ mice were significantly more sensitive to insulin than WT mice (Figure 1G). Insulin tolerance test (ITT) confirmed that DC^hBcl-2^ mice displayed enhanced insulino-sensitivity compared to WT mice (Figure 1H, I).

We further investigated the DIO resistant phenotype of DC^hBcl-2^ mice evaluating their daily energy expenditure. We monitored the daily food intake and observed no differences between WT and DC^hBcl-2^ mice upon HFD (Figure 1J). In parallel, we evaluated the intestinal absorptive capacity looking at loss of energy in the feces by bomb calorimetry. We observed no significant differences in fecal kilocalories excreted per g of feces per day (kcal/g/d) in both groups of mice with no differences observed in term of transit time (Figure 1K – figure supplement 1B) and feces production (figure supplement 1C). Indirect gas calorimetry and locomotor activity assessment indicated that there were no changes between WT and DC^hBcl-2^ mice upon HFD (figure supplement 1D, G).

All these results suggested that differences in weight gain, adiposity and insulin-sensitivity in HFD-fed WT and DC^hBcl-2^ mice occurred despite similar food intake, caloric intestinal absorptive capacity and energy expenditure.

DC^hBcl-2^ mice are resistant to HFD induced obesity and associated metabolic alterations. Body weight (A) and fat mass (C) monitoring of WT and DC^hBcl-2^ at day 0, 12 and 24 weeks after starting high fat diet (HFD). (B) Abdominal photographs of representative WT and DC^hBcl-2^ mice HFD-fed for 24 weeks. (D) Average adipocyte diameter in the sub-cutaneous adipose tissue of WT and DC^hBcl-2^ HFD-fed for 24 weeks (N=7 to 9 mice per group). (E) Blood glucose (g/L) and (F) insulin (ng/mL) levels during oral glucose tolerance test (OGTT) after 13 weeks of HFD (N=10 to 14 mice per group). (G) Insulin resistance index (HOMA-IR) after 13 weeks of HFD (N=10 to 14 mice per group). (H) Blood glucose level (g/L) during an insulin tolerance test (ITT) after 14 weeks of HFD (N=10 to 14 mice per group). (I) Area under the curve of the glucose profile during the ITT (N=10 to 14 mice per group). Mean of daily food intake (J) and fecal energy (K) monitored for one week after 12-weeks of HFD. Data are presented as median for dot plots and mean ± SD for others.

### Intestinal barrier integrity in DC^hBcl-2^ mice is associated with increased tolerogenic DCs

Since impaired intestinal barrier function (IBF) has been involved in the development of obesity and associated metabolic alterations, we investigated how this function was affected by the HFD in WT and DC^hBcl-2^ mice.

We evaluated the intestinal paracellular permeability by monitoring the appearance of fluorescence in the blood following oral administration of fluorescein isothiocyanate (FITC)-dextran. DC^hBcl-2^ mice displayed a lower paracellular intestinal permeability compared to WT mice (Figure 2A). This was associated with lower levels of fecal albumin (Figure 2B), used as a marker for gut vascular barrier leakage (Lirui Wang et al. 2015). We determined how these differences in term of intestinal permeability relate to the overall intestinal inflammatory tone. We quantified fecal lipocalin2 (Lcn2), known as an early biomarker for intestinal inflammation (Chassaing et al. 2012). DC^hBcl-2^ mice displayed lower levels of lipocalin 2 compared to WT mice (Figure 2C). Fecal Lcn2 levels observed in WT mice were comparable to a low-grade inflammation as previously shown in HFD-mediated experimental models (Natividad et al. 2018).

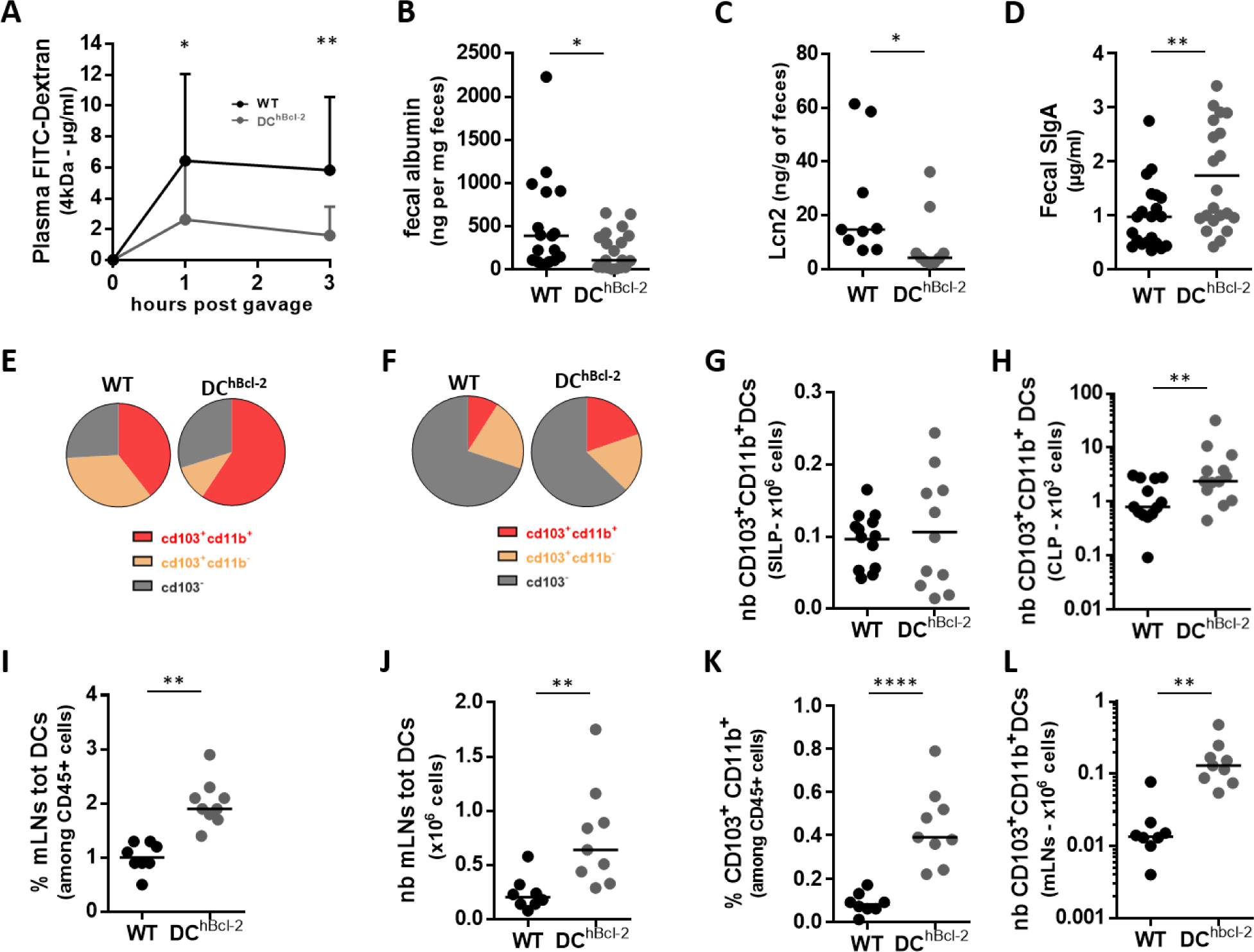

IBF also comprises fecal SIgA that counteract several antigens to access the intestinal wall (bacteria, food etc.) (Mantis, Rol, and Corthésy 2011). Fecal SIgA titers were 1.8 times higher in DC^hBcl-2^ than in WT mice (Figure 2D).

All these results suggested that DC^hBcl-2^ mice displayed enhanced IBF compared to WT mice after 24 weeks of HFD.

We further assessed how manipulating DCs lifespan impacted intestinal DCs populations. We first looked at global markers and subsets of DCs by flow cytometry in WT and DC^hBcl-2^ intestines (figure supplement 2A). We observed no significant differences both in the proportion and in the total numbers of DCs (totDCs) in both small intestine lamina propria (SILP) and colon lamina propria (CLP) (figure supplement 2C, F). Three subsets of intestinal conventional DCs (cDCs) have been described, depending on the CD103 and CD11b surface markers (Merad et al. 2013; Bekiaris, Persson, and Agace 2014). Looking deeper in cDCs subpopulations, we found that both the SILP and the CLP CD103^+^ CD11b^+^ cDCs were significantly increased in proportion of total DCs in DC^hBcl-2^ compared to WT mice (Figure 2E, F). Despite no difference was observed in the SILP, total numbers of CD103^+^ CD11b^+^ cDCs (Figure 2G) in the CLP were 3-fold increase in DC^hBcl-2^ mice relatively to WT mice (Figure 2H). Those results demonstrated that the *hbcl2* insertion strongly enhanced the tolerogenic CD103^+^ CD11b^+^ DCs subpopulation.

One important feature of DCs after antigenic stimuli is their ability to migrate from the intestinal lamina propria to the mesenteric lymph nodes (mLNs). We therefore characterized DCs populations in the mLNs of WT and DC^hBcl-2^ mice after 24 weeks of HFD. We observed a marked increase in total DCs (totDCs) in both, percentage and total numbers (Figure 2I, J) in the mLNs of DC^hBcl-2^ mice compared to WT mice. The CD103^+^ CD11b^+^ cDCs subset in DC^hBcl-2^ mice was even more increased in this compartment, representing four times more in proportion than in WT mice and ten times more in total numbers (Figure 2K, L).

These observations highlighted that the maintenance of DC^hBcl-2^ mice intestinal barrier function is associated with an increase in the tolerogenic CD103^+^ CD11b^+^ cDC subset.

Maintenance of the IBF is associated with a strong increase in tolerogenic DCs. All data are representative of mice fed a HFD for 24 weeks. (A) Plasma levels of dextran-FITC at 1 and 3 hours post-oral gavage (600 mg/kg body weight) (N=10 to 14 mice per group). Albumin (B), Lcn2 (C) and secreted immunoglobulin-A (SIgA) (D) levels in the feces determined by ELISA. (E) (F) Mean proportions of CD103^+^ CD11b^+^, CD103^+^ CD11b^-^, CD103^-^ among total DCs in the SILP (E) or in the CLP (F). (G)(H) Total numbers of CD103^+^ CD11b^+^ DCs in the SILP (G) or in the CLP (H). Proportions (I) and total numbers (J) of total DCs among CD45^+^ cells isolated from the mLNs. (K) Proportions and (L) total numbers of CD103^+^ CD11b^+^ DCs in the mLNs. Data are presented as mean for circle graphs, median for dot plots and mean ± SD for others.

### *hBcl2* transgene promotes DCs tolerogenic properties with enhanced RALDH activity

Previous research have demonstrated that upon DC maturation and notably after bacterial stimulation, Bcl2 expression (gene and protein) was downregulated (Granucci et al. 2001; Nopora and Brocker 2002). To understand how the *hBcl2* transgene insertion modulates more particularly the tolerogenic DCs population we performed global transcriptomic analysis on sorted DCs. We focused on DCs from the mLNs where adaptive immunity priming occurs. A first screen of mLNs DCs subpopulations indicated that the *hBcl2* transgene was significantly more expressed in the CD103^+^ DCs than in CD103^-^ DCs regardless of the diet (Figure 3A). Among the CD103^+^ DCs, CD11b^+^ DCs were the most enriched in transgenic compared to WT mice (figure supplement 3A). We assessed the effect of *hBcl2* on CD103^+^CD11b^+^ sorted DCs, which appeared the most affected by the transgene insertion in the mLNs of both groups of mice. To avoid any other environmental effect that could synergize with the *hBcl2* insertion we performed their global gene expression analysis in mice before starting the HFD. Principal component analysis of CD103^+^ CD11b^+^ gene sets discriminated the sample genotypes on the high axe percentages (69.5% and 7.1% for x and y axes, respectively) confirming the major impact of the transgene expression in this particular cDCs subtype (Figure 3B).

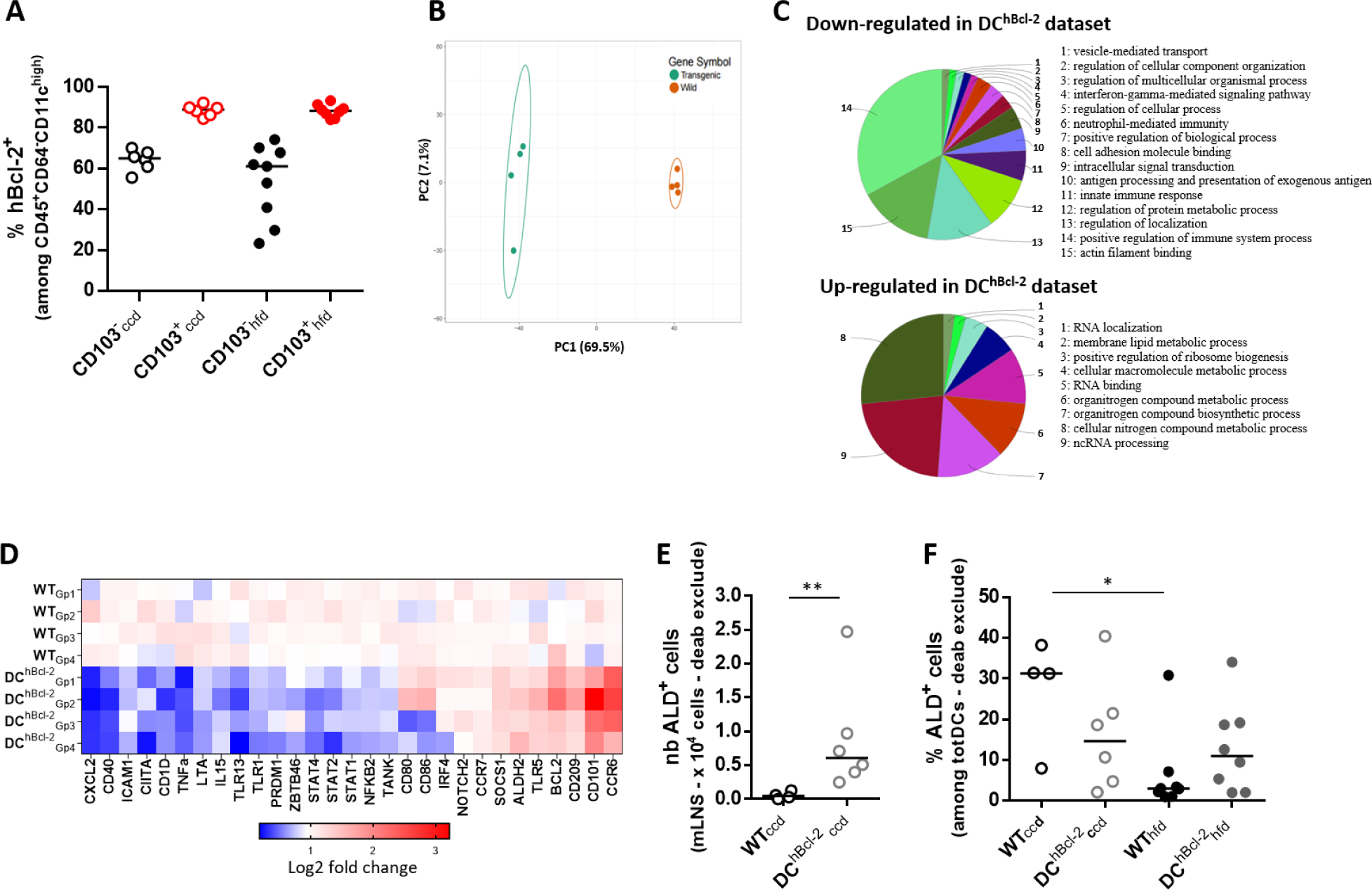

Differential expression of genes calculated by Student t-test showed that 2774 genes discriminate DC^hBcl-2^ and WT samples with a p value < 0.05 corrected by Benjamini-Hochberg for false discovery rate (FDR) (Benjamini and Hochberg 1995).

Looking deeper into pathways up- or down-modulated in the two datasets, we noticed a global down regulation of immune-related pathways in DCs sorted from DC^hBcl-2^ mice (Figure 3C). Expression of genes related to the maturation or activation status of DCs were downregulated in DC^hBcl-2^ CD103^+^ CD11b^+^ DCs (CXCL2, CD40, CIITA, CD1D) (Figure 3D). These results were confirmed looking at predictive signaling pathways involved in DCs maturation using Ingenuity Pathway Analysis (figure supplement 3B). We observed that the human anti-apoptotic factor Bcl-2 (hBcl-2) may prevent DCs to acquire antigen sensing through TLRs 2/3/4/9, antigen-presenting properties through MHC class II/I or cell adhesion markers as the Intercellular Adhesion Molecule 1 (ICAM1) as well as markers for co-stimulation of adaptive immune cells (CD40/ CD86) (figure supplement 3B). Functionally DCs immaturity relates to their inability to mount pro-inflammatory responses after stimulation (Mahnke et al. 2002). The immature/inactive status of transgenic sorted-DCs was in line with a down-
regulation of immune-related inflammatory signaling pathways as nuclear factor kappa-light-chain-enhancer of activated B cells pathway (NF-kB, TANK, NFKB2) and signal transducer and activator of transcription pathways (STAT1-2-4) (Figure 3D – figure supplement 3B). This global down-regulation of pro-inflammatory responses was associated with a decreased capacity for pro-inflammatory cytokines release such as TNFa, IFNg and IL-15 (Figure 3D – figure supplement 3B). All these results strongly suggest that *hBcl2* transgene prompted CD103^+^CD11b^+^ DCs to keep an immature phenotype altering their capacity to elicit the pro-inflammatory immune responses.

Immature immune stage in DCs has been associated with increased tolerogenic capacity (Mahnke et al. 2002; Tisch 2010). Expression levels of gene sustaining tolerogenic functions such as CD101, SOCS1 as well as ALDH2 were upregulated in DC^hBcl-2^ CD103^+^ CD11b^+^ DCs compared to WT (Figure 3D).

Since tolerogenic ability of mucosal DCs has been related to their capacity to process vitamin A into retinoic acid through enzymatic activity of retinaldehyde dehydrogenases (RALDHs) (Cassani et al. 2012), we next analyzed this function in CD103^+^ CD11b^+^ DCs isolated from mLNs of both group of mice maintained under control chow diet (CCD). We performed an immuno-staining of RALDH activity and analyzed by flow cytometry this enzymatic function (figure supplement 3C). We observed enhanced RALDH activity (ALD^+^ cells) in CD103^+^ CD11b^+^ DCs subpopulation isolated from mLNs of DC^hBcl-2^ mice. ALD^+^ cells represented a 3-fold increase in percentage relatively to WT mice that resulted in a 13-fold increase in total ALD^+^ cell number among CD103^+^ CD11b^+^ cells (Figure 3E, F).

Considering the strong increase of RALDH activity in the mLNs of DC^hBcl-2^ mice in CCD, we wonder how this DC function was impacted upon HFD. We looked at mLNs DCs RALDH activity and observed no impact of the diet on the RALDH activity in both groups, DC^hBcl-2^ mice significantly maintaining their higher rate of RALDH activity (Figure 3E, F). Studies have highlighted the importance of DCs RALDH activity in mouse models of colitis and in inflammatory bowel disease patients (Laffont, Siddiqui, and Powrie 2010; Magnusson et al. 2016). In those studies, they demonstrated that upon inflammation tolerogenic DCs were losing their RALDH activity. This participated to establish a pro-inflammatory immune environment promoting the subsequent loop of chronical inflammation. Despite different levels of inflammation between inflammatory bowel diseases and metabolic diseases, concordant immune dysfunctions related to intestinal barrier leakage have been observed (Winer et al. 2016). We wondered how HFD would impact the intestinal RALDH DCs function in both group of mice. Despite we observed no difference in the small intestine lamina propria (data not shown), DCs RALDH activity was decreased in the colon lamina propria (CLP) of WT mice in HFD condition compared to CCD (Figure 3F). On the contrary, DC^hBcl-2^ mice maintained their RALDH activity (Figure 3F – figure supplement 3G).

Those results indicated that modulating DCs through *hBcl2* insertion promotes tolerogenic DCs with increased capacity to process vitamin A through RALDH activity.

*hBcl2* transgene modulates tolerogenic DCs function that impact RALDH activity. (A) Proportion of hBcl-2^+^ DCs among CD103^+^ and CD103^-^ subpopulations in the mLNs after 24 weeks of CCD or HFD. (B) (C) (D) Microarray gene expression analysis of sorted CD103^+^ CD11b^+^ DCs from mLNS of WT and DC^hBcl-2^ before starting the diet (B) Principal component analysis showing separation of sample groups (C) Biological enrichment and annotation of pathways down-regulated and up-regulated in DC^hBcl-2^ dataset using ClueGo plugin. (D) Heatmap of log 2-fold-change value of key gene expression related to DC maturation or activation and DCs tolerogenic markers. (E) Total numbers of aldefluor positive cells (ALD^+^) in CD11b^+^ CD103^+^ cDCs subpopulation in the mLNs of mice before starting the diet. (F) Total numbers of ALD^+^ cells among total DCs in the CLP of mice after 24 weeks of CCD or HFD. Data are presented as median for dot plots.

### Tolerogenic DCs strongly impact colonic adaptive immunity in the context of DIO

The important role of DCs in shaping the appropriate immune responses through adaptive immune system activation has been widely documented at steady state as well as in many inflammatory conditions (Coombes and Powrie 2008; Bekiaris, Persson, and Agace 2014). After 24 weeks of HFD, we observed no significant differences in proportions and total numbers of T and B lymphocytes in the mLNs of both groups of mice (figure supplement 4A, C – data not shown). Focusing on T helper subsets, we detected higher proportions and total numbers of CD4^+^ IL17^+^ cells (Th17) as well as higher total numbers of CD4^+^ Foxp3^+^ (Treg) in the mLNs of DC^hBcl-2^ compared to WT mice (Figure 4A, 4B and figure supplement 4D, 4E). In the same compartment, we also observed higher proportion and total numbers of CD19^-^ sIgA^+^ plasmablasts in DC^hBcl-2^ mice (Figure 4C, D).

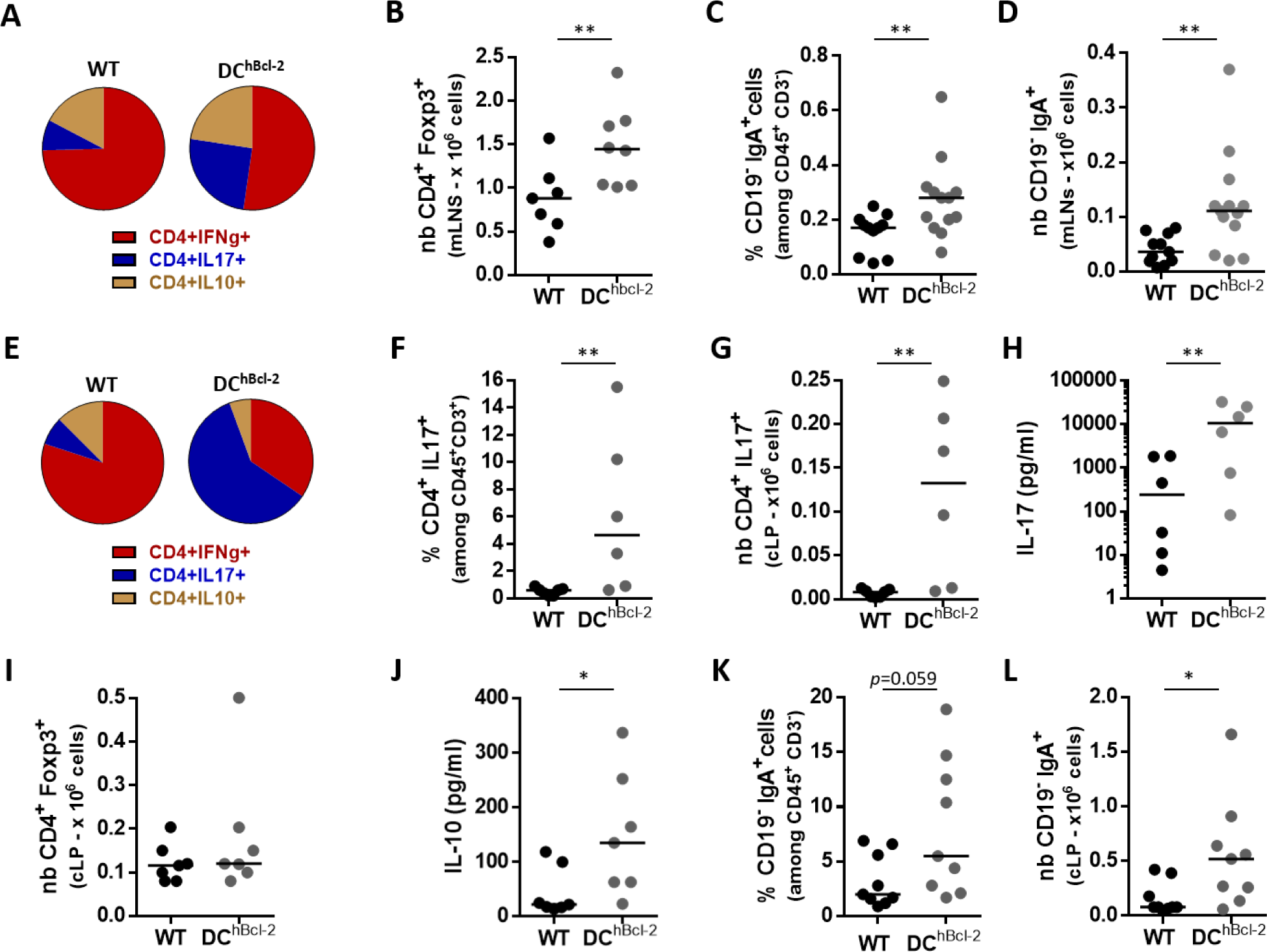

Altogether these results demonstrated that higher number of tolerogenic CD103^+^ CD11b^+^ cDCs were associated with enhanced priming of Th17, Treg and sIgA^+^ B cell responses in DC^hBcl-2^ after 24 weeks of HFD.

We next assessed where those adaptive immune responses established throughout the intestinal compartment. We observed no significant differences in CD4^+^ T cell responses or sIgA^+^ B cell responses in the SILP compartment (figure supplement 4F – 4G). Conversely, we observed a marked and significant skewing toward Th17 responses in the CLP of DC^hBcl-2^ mice upon HFD. While Th1 cells were the predominant T cell population in the CLP of WT mice, Th17 cells represented 60% of the total CD4^+^ T cells in DC^hBcl-2^ mice (Figure 4E). DC^hBcl-2^ mice Th17 colonic responses displayed a 4-fold increase in percentage and a 10-fold increase in total number relatively to WT mice (Figure 4F, G). Cellular ex vivo experiments with a non-specific TCR stimulation confirmed that colonic T cells isolated from DC^hBcl-2^ mice displayed higher capacity to secrete IL-17 than those from WT mice (Figure 4H). Despite equivalent levels of CD4^+^ Foxp3^+^ Treg cells, ex vivo stimulated colonic T cells isolated from DC^hBcl-2^ mice secreted higher levels of IL-10 (Figure 4I, J). Th17 and Treg responses that established in the colon of DC^hBcl-2^ mice were associated with a significant increase in proportion and total number of CD19^-^ sIgA^+^ plasmablasts (Figure 4K, L).

Altogether those results suggested that upon HFD, increased tolerogenic DCs strongly impact the colonic immunity through enhanced Th17, Treg and sIgA^+^ B cell responses.

HFD-fed DC^hBcl-2^ mice showed enhanced Treg Th17 and sIgA^+^ B cells that established in the colon. All data are representative of mice fed a HFD for 24 weeks. (A) (E) Circle graphs representing the mean proportions of IFNg-producing, IL-17-producing, IL-10-producing CD4^+^ T cells in the mLNs (A) or in the CLP (E) after intracellular staining of cytokines. (B) (I) Total numbers of CD4^+^ Foxp3^+^ T lymphocytes in the mLNs or in the CLP (I). (C) (K) Proportions of CD19^-^ IgA^+^ plasmablasts in the mLNs (C) or in the CLP (K). (D) (L) Total numbers of CD19^-^ IgA^+^ plasmablasts in the mLNs (D) or the CLP (L). (F) Proportions and total numbers (G) of IL-17-producing CD4^+^ T cells in the CLP after intracellular staining of cytokines. (H) IL-17 and IL-10 (J) secretion in the supernatants of ex-vivo anti-CD3/CD28 stimulated CLP cells for 72h. Data are presented as mean for circle graphs or median for dot plots and.

### DC^hBcl-2^ intestinal microbiota displays lower inflammatory signatures after DIO

The immune system strongly influences intestinal microbiota composition that, in turn, is a strong determinant of the metabolic response to HFD.(Belkaid and Hand 2014; Le Roy et al. 2013) We wondered how the strong colonic immunological differences that we observed could impact the intestinal microbiota in both groups of mice. We analyzed the fecal microbiota of WT and DC^hBcl-2^ mice by 16S rRNA gene sequencing before and after HFD challenge. 16S rRNA gene analyses first indicated that the fecal microbiota of WT and DC^hBcl-2^ mice were not distinguishable before starting the diet (Figure 5A, B). Conversely, we observed marked differences in WT and DC^hBcl-2^ microbiota after twelve weeks of HFD (Figure 5C, D). Linear discriminant analysis (LDA) effect size (LEfSe) revealed that specific taxa were enriched in either the microbiota of WT or the DC^hBcl-2^ mice. WT mice harboured enrichment of the *Epsilonproteobacteria* family members whereas the *Mogibacteriaceae* members appeared increased in DC^hBcl-2^ mice (Figure 5E, F). Those results demonstrated that upon HFD, microbiota composition has differentially shifted upon HFD in the two groups of mice.

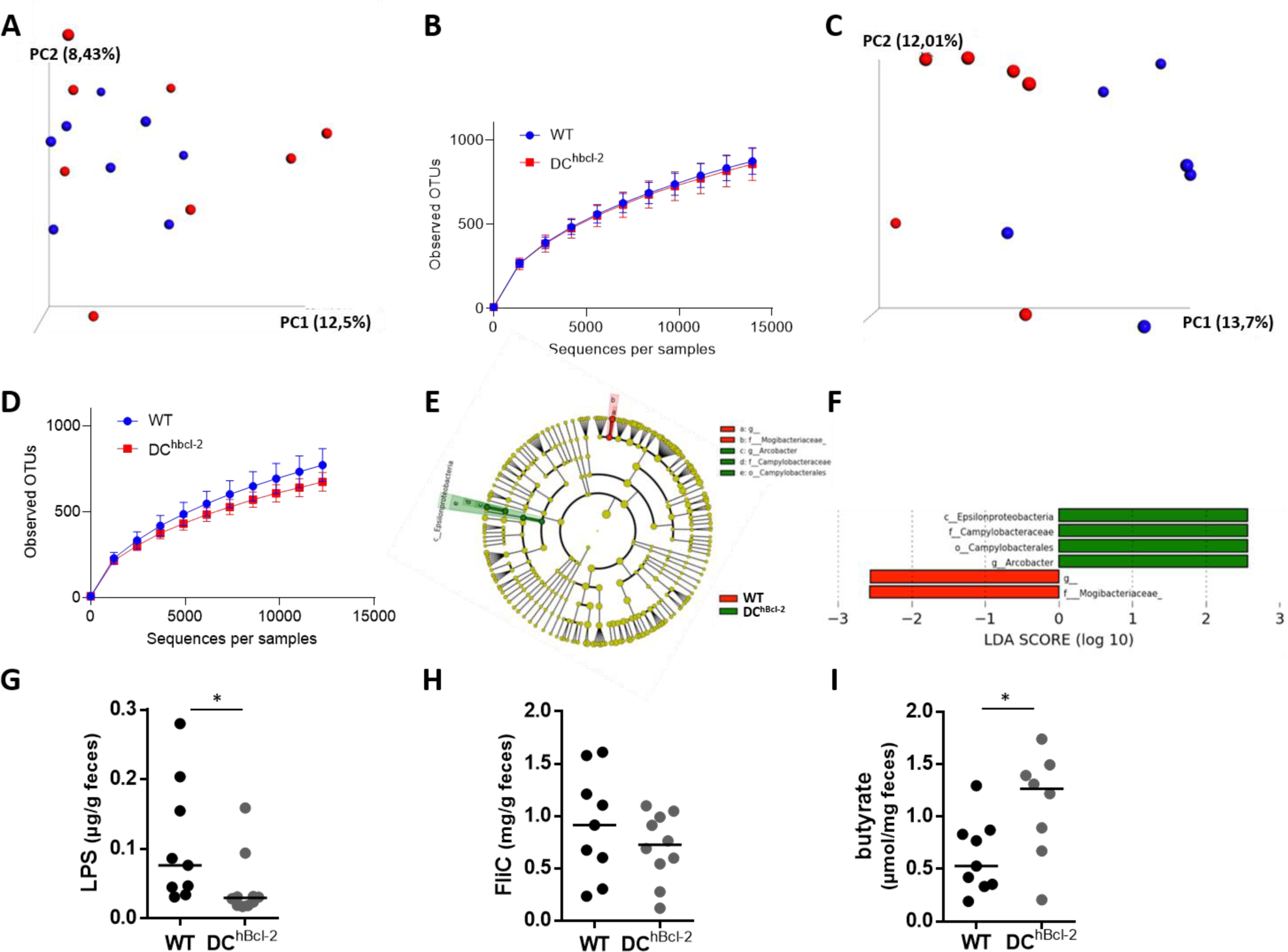

*Epsilonproteobacteria* are members of Gram-negative bacteria. Their motility as well as their lipopolysaccharides (LPS), major components of their outer membrane, could trigger pro-inflammatory immunoreactivity. We assessed the immunogenic properties of each type of microbiota through the quantification of fecal bioactive LPS and fecal bioactive flagellin (FliC) using the system reporter cell lines for murine Toll like Receptor (TLR) type 4 and type 5 respectively. Upon HFD, DC^hBbcl-2^ mice displayed lower amount of both fecal bioactive LPS and FliC and this was more particularly significant for the bioactive LPS (Figure 5G, H).

Although it has been widely demonstrated that DCs sense microbiota-derived signals through PRRs like TLRs, other sets of PRR-independent signals can orientate DC function. Bacterial fermentation products have been shown to participate in the immunoregulatory function of DCs, contributing to intestinal immune tolerance and maintenance of intestinal homeostasis (Zhao and Elson 2018). We quantified fecal short chain fatty acids (SCFA) using gas–liquid chromatography. Despite comparable amount of total fecal SCFA concentration (figure supplement 5C), DC^hBcl-2^ harbored particular SCFA profiles with a marked enrichment in fecal butyrate concentration, representing 2,3-fold increase relatively to WT mice (Figure 5I).

Altogether those results demonstrated that under HFD, DC^hBcl-2^ microbiota behave differently in terms of bacterial composition and functions leading to less immunogenicity as well as sustaining immune tolerance.

HFD-fed DC^hBcl-2^ mice shape a gut microbiota characterized by lower inflammatory signatures. (A-D) Principal component analysis (PCA) of the unweighted UniFrac distance matrix (A, C) and alpha diversity assessment (B, D) of fecal WT and DC^hBcl-2^ microbiota at baseline (A, B) and after 12 weeks of HFD (C, D). (E, F) LEfSe (LDA Effect Size) was used to investigate bacterial members that drive the differences between the fecal microbiota of WT and DC^hBcl-2^ mice. (E) Taxonomic cladogram obtained from LEfSe analysis. Red, taxa significantly more abundant in WT mice; green, taxa significantly more abundant in DC^hBcl-2^ mice. (F) LDA scores for the differentially altered taxa.

Green, taxa significantly more abundant in WT mice; red, taxa significantly more abundant in DC^hBcl-2^ mice. Only taxa meeting an LDA significance threshold > 2.0 are presented. (G, H) Fecal LPS and FliC levels in WT and DC^hBcl-2^ mice assessed by HEK reporter cell lines. (I) Fecal butyrate concentrations in WT and DC^hBcl-2^ mice. Data are represented as median for dot plots.

### DC^hBcl-2^ intestinal microbiota drives resistance to HFD-induced metabolic alterations

To unravel the respective role of WT and DC^hBcl-2^ microbiota in modulating their metabolic phenotype we first compared HFD-treated co-housed WT (WT CoH) and DC^hBcl-2^ (DC^h^ CoH) mice versus single housed WT or single housed DC^hBcl-2^ mice. Looking at body mass composition after 24 weeks of HFD we observed that cohousing transmitted the DIO-resistant phenotype to WT mice (figure supplement 6A – 6C).

To assess whether the microbiota was driving the HFD-resistant DC^hBcl-2^ phenotype, we transferred fecal microbiota from both groups of mice into germ-free (GF) recipients. We colonized 8 weeks old GF recipients with the microbiota of WT and DC^hBcl-2^ mice that were previously fed a HFD during 24 weeks (figure supplement 6D). After 24 weeks of diet, DC^hBcl-2^ microbiota recipient mice (i.e. FT-DC^hBcl-2^) gained significantly less weight than WT microbiota recipient ones (i.e. FT-WT) (39,2g ± 3,9g and 44,2g ± 3,9g respectively) (Figure 6A). Looking more precisely at body mass composition, we observed that mice developed the same lean mass (figure supplement 6E), but that FT-DC^hBcl-2^ mice displayed significantly less adiposity than FT-WT mice (10,7g ± 2,3g and 15,6g ± 3,2g respectively) (Figure 6B).

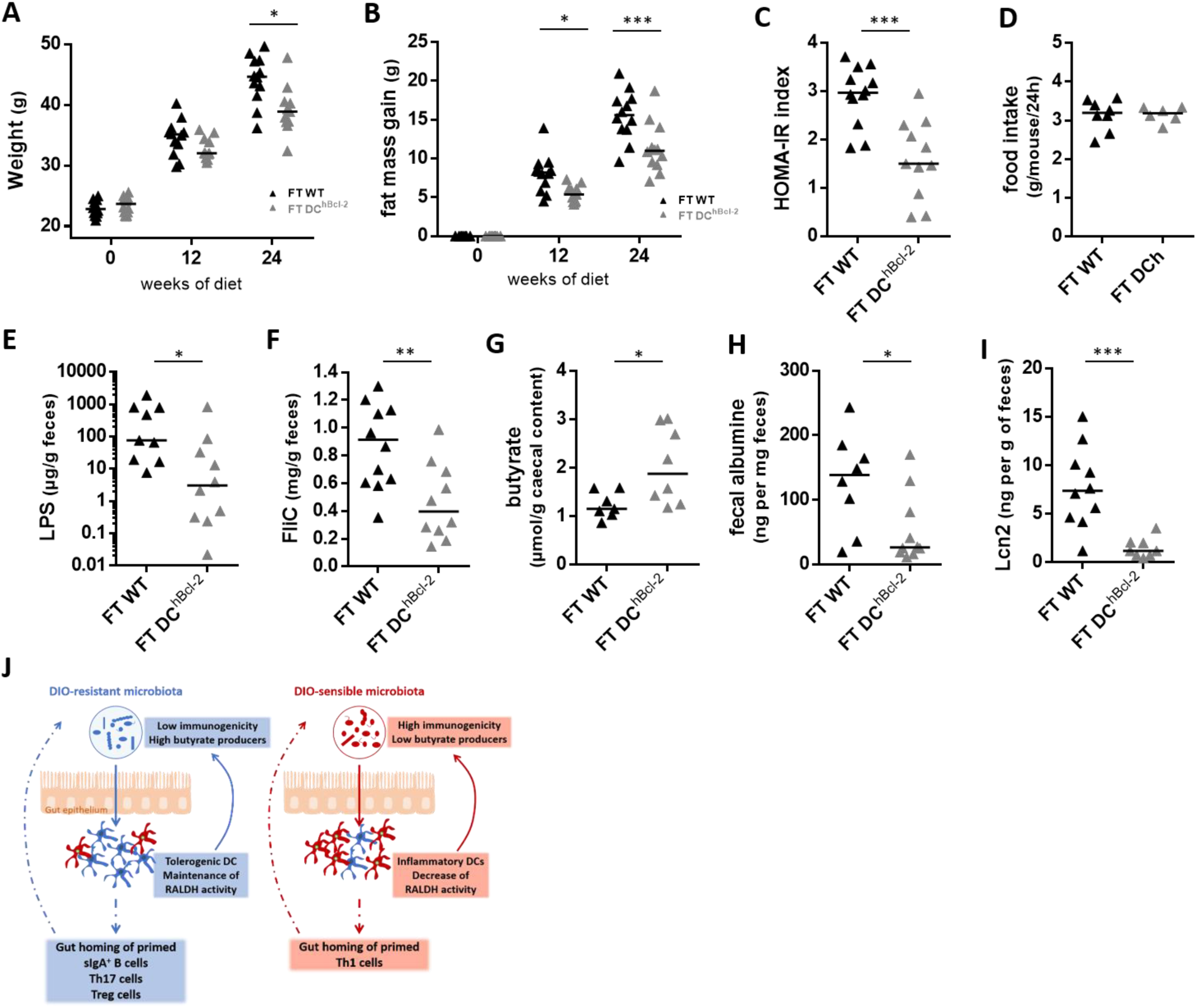

We further investigated how glucose metabolism was impacted after microbiota transplantation and observed marked differences in insulin sensitivity. FT-DC^hBcl-2^ recipient mice had a significantly lower HOMA-IR index than FT-WT mice (1,6 ± 0,8 and 2,9 ± 0,6 respectively) (Figure 6C). As monitored in donors, we never observed any variations in food intake in the recipient groups (Figure 6D).

Those results were in line with cohousing experiments and demonstrated that DC^hBcl-2^ microbiota by itself was able to drive the HFD-resistant phenotype. This overall demonstrated that DC^hBcl-2^ tolerogenic DCs are shaping a transmissible DIO-protective intestinal microbiota.

We next assessed whether the immunogenic properties of each type of microbiota have been transmitted through fecal transplantation. FT-DC^hBcl-2^ recipient mice displayed a significant lower amount of both fecal bioactive LPS (Figure 6E), and fecal bioactive FliC (Figure 6F). We quantified caecal SCFA and observed that FT-DC^hBcl-2^ recipients harbored enrichment in caecal butyrate concentration, representing 1,6-fold increase compared to FT-WT recipients (Figure 6G). We wondered if these discrepant microbial properties transferred from donors to recipient mice have impacted intestinal permeability looking at fecal albumin content. We observed a 3-fold increase in fecal albumin content in FT-WT mice relatively to FT-DC^hBcl-2^ mice (Figure 6H). After 24 weeks of HFD FT-DC^hBcl-2^ recipients developed less intestinal inflammation than FT-WT recipients as demonstrated by the lower level of fecal lipocalin2 with respective means and SD of 1,5 ± 4,2 and 7,8 ± 1,0 ng/g of feces (Figure 6I).

Altogether those results demonstrated that an increase in tolerogenic DCs is associated with a DIO-resistant microbiota that is sufficient to drive by itself the DIO resistant phenotype by cohousing or after fecal microbial transfer into germ-free recipients. This transmissible DIO-resistant microbiota is characterized by less immunogenicity, enhanced butyrate producing capability related with decreased intestinal inflammatory tone.

In summary, our data revealed how enhancing tolerance through targeting DC survival pathway can promote DIO resistance, with a central role played by the intestinal microbiota. The adaptive immune responses that established in the intestinal compartment under DCs tolerogenic pressure may participate to counteract the HFD-mediated increase in microbial immunogenicity and may favour immunoregulatory microbial functions through butyrate production (Figure 6J). Hence, in the context of metabolic syndrome, assessing the tolerogenic function of DCs in patients may provide new insight for diagnostic and therapeutic approaches.

DC^hBcl-2^ gut microbiota triggers the DIO-resistant phenotype. (A) Body weight monitoring of WT-microbiota recipients (FT-WT) and DC^hBcl-2^ -microbiota recipients (FT-DC^hBcl-2^) at day 0, 12 and 24 weeks after both fecal microbial transplant (FT) and starting the HFD. (B) Fat mass gain at day 0, 12 and 24 weeks after both FT and HFD. (C) Insulin resistance index (HOMA-IR) after 12 weeks of both FT and HFD. (D) Mean of daily food intake of individually-housed mice monitored for one week after 12 weeks of both FT and HFD. Fecal bioactive LPS (E) and FliC (F) load after 24 weeks of both FT and HFD. (G) Caecal butyrate concentration after 24 weeks of both FT and HFD. Albumin (H) and Lcn2 (I) levels in the feces after 24 weeks of both FT and HFD. (J) Scheme representing how tolerogenic DCs sustain the DIO-resistant microbiota characterized by lower immunogenicity and enhanced butyrate production. Data are presented as median for dot plots

## Discussion

Bcl-2-regulated apoptosis pathway has been shown to act as a molecular regulator of both DC lifespan and immunogenicity (Hou and Van Parijs 2004; Nopora and Brocker 2002; Gautier et al. 2008; Gautier Emmanuel L. et al. 2009). The functional importance of this survival pathway tested in vivo, in the context of acute exposure to non-lethal doses of LPS, revealed that Bcl-2 regulate accumulation of DCs associated with enhanced T cell activation which in turn enables resistance to lethal septic shock in mice (Gautier et al. 2008). Here, we questioned what could be the impact of such DC-mediated Th polarization in the context of HFD-induced metabolic endotoxemia, where LPS is playing a central role in driving the deleterious metabolic effect (Cani et al. 2007a; 2008; X. Wang et al. 2014). The DC^hBcl-2^ DIO-resistant phenotype was indeed characterized by healthier indexes of intestinal barrier function together with a lower inflammatory tone. Characterization of the intestinal immune responses demonstrated a marked enrichment toward CD103^+^ CD11b^+^ cDCs, especially in the colon of DC^hBcl-2^ mice. This particular cDCs subpopulation has been shown to induce the differentiation of Th17 cells in the gut at steady state (Persson et al. 2013), and we indeed observed a strong colonic Th17 polarization in the intestinal draining lymph nodes as well as in the colon lamina propria of DC^hBcl-2^ compared to WT mice. Previous studies have shown that DIO triggers an increase of intestinal Th1 immune response associated with a decrease of intestinal Th17 response (Luck et al. 2015; Garidou et al. 2015; Hong et al. 2017). In Garidou *et al*. and Hong *et al*., the authors even demonstrated the important role of Th17 cells in mediating DIO resistance. Our data are in line with what has been published reinforcing the hypothesis that intestinal Th17 responses play a major role in counteracting DIO and metabolic alterations.

The mechanism by which intestinal Th17 responses are decreased under DIO is still questioned. One possible explanation could be a lack for a proper antigen stimulation of T cells by antigen presenting cells (Garidou et al. 2015; Hong et al. 2017). Since DCs are mainly involved in this process, we investigated how the hBcl-2-targeted CD103^+^ CD11b^+^ cDCs subpopulation may have induced Th17 polarization. We found out that RALDH tolerogenic DCs function, converting the vitamin A into retinoic acid, is increased in DC^hBcl-2^ mice relatively to WT mice. The importance of such DCs tolerogenic activity has been previously demonstrated at steady state. DC-derived RALDH retinoic acid production has been shown to regulate adaptive immune responses within the intestine, thereby controlling functional T and B-cell differentiation and directing their migration toward the intestine (Mora et al. 2006; Iwata et al. 2004; Mucida et al. 2007; Coombes et al. 2007). In a context of inflammation, and more precisely in the colon of ulcerative colitis patients, the RALDH DCs activity is impaired (Magnusson et al. 2016). Despite a lack of evidence relating this decreased function to disease progression, retinoic acid treatment in both human biopsies and animal models of ulcerative colitis decreases the inflammation especially through the induction of T regulatory responses (Bai et al. 2009). In the context of HFD, vitamin A deficient diet worsens the metabolic phenotype and is associated with a more severe decrease of intestinal Th17 cells (Hong et al. 2017). The overall enhanced tolerogenic DC activity in the intestinal compartment including draining lymph nodes could explain the discrepant intestinal adaptive immunity that established in DC^hBcl-2^ and WT mice. Indeed Treg, Th17 and sIgA^+^ B cells, all increased in DC^hBcl-2^ compared to WT mice, are major components of intestinal homeostasis (Li Wang, Zhu, and Qin 2019).

Several researches demonstrated how the adaptive immunity is impacting the systemic metabolism through intestinal microbiota modulations (Garidou et al. 2015; Hong et al. 2017; Petersen et al. 2019). Th17-mediated DIO-resistance involves their ability to control microbiota composition (Garidou et al. 2015; Hong et al. 2017). We demonstrated here, through cohousing and fecal microbiota transplantation approaches, that DC^hBcl-2^ microbiota is sufficient to transmit the DIO-resistance phenotype. Analysis of fecal microbial composition revealed that WT mice depicted an enrichment toward the *Epsilonproteobacteria* members under HFD. Those Gram-negative bacteria represent an important source of immunogenic LPS, known to trigger metabolic endotoxemia as previously demonstrated (Cani et al. 2008; 2007b). The respective microbial immunogenic property was confirmed by looking at the fecal load of bioactive LPS, which was increased in the WT fecal microbiota compared to DC^hBcl-2^. This particular immunogenic trait of DIO-sensible WT microbiota was transmitted to the recipients, suggesting that such discrepant immunogenic load of fecal LPS plays a role in the different phenotypes resulting from DIO treatment.

Another interesting bacterial component increased in DIO-resistant DC^hBcl-2^ mice compared to WT mice is butyrate, a SCFA involved in many metabolic processes promoting host fitness and shaping the intestinal immune system (Gao et al. 2009; De Vadder et al. 2014; Parada Venegas et al. 2019). Although butyrate has been related to impact feeding behaviour and/or energy expenditure, we never noticed any differences in term of food intake nor any other parameters using metabolic cages. Instead of a direct action on energy balance, the increased butyrate content could sustain the immunoregulatory responses that were enhanced in DIO-resistant DC^hBcl-2^ mice. Several researches have indeed highlighted the important role of butyrate in downregulating the expression of pro-inflammatory immune responses as well as promoting immunoregulatory ones (Arpaia et al. 2013; Li et al. 2018). DC^hBcl-2^ mice harboured increased polarization of Treg cells in their gut draining lymph nodes and HFD-treated DC^hBcl-2^ colonic Treg displayed enhanced capacity to produce IL-10 compared to WT mice. Another potent mechanism of butyrate-mediated immunoregulatory process could have directly impacted the local tolerogenic capacity of DCs which in turn have been shown to sustain Treg activity (Singh et al. 2014). It has been more particularly demonstrated that butyrate-conditioned human DCs are able to prime Treg cells through the induction of RALDH function (Kaisar et al. 2017). This latter observation overall reinforces the importance of RALDH tolerogenic DCs function, and demonstrates how microbial-derived metabolites may sustain these DCs mediated immunoregulatory activities. Furthermore, with the observation that butyrate was also increased in the FT-DC^hBcl-2^ recipients, our results strongly suggest that such discrepant microbiota compositions and functions can trigger metabolic phenotype even in the absence of the transgene.

## Materials and Methods

### Animal experimentation

Transgenic male mice on the C57BL/6J background expressing hBcl-2 under the murine CD11c promoter (DC^hBcl-2^) were obtained as previously described(Gautier et al. 2008). Littermates at birth until weaning, heterozygous DC^hBcl-2^ and wild type (WT) controls were either cohoused or single housed depending on their genotype in Individually Ventilated Cages (IVC). Mice were fed either a control chow diet (CCD) (E157451-347, ssniff Spezialdiätten GmbH, Soest, Germany) or a high-fat diet (HFD) (60 % fat and 20 % carbohydrates (kcal/kg), E15742-347, ssniff Spezialdiätten GmbH, Soest, Germany) starting at 8-weeks old of age for 24 weeks. Mice had free access to food and water. Body composition was assessed by using 7.5 MHz time domain-nuclear magnetic resonance (TD-NMR) (LF90II minispec, Bruker, Rheinstetten, Germany) at 0, 12 and 24 weeks of diet. The mice were killed by cervical dislocation and organs were collected, frozen in liquid nitrogen or processed for single cells preparation. All procedures involving mice were carried out according to the Guide for the Care and Use of Laboratory Animals published by the European Commission Directive 86/609/EEC. All animal studies were approved by the regional veterinary services of the Paris police headquarters (agreements no. 75-751320) and by the Biological Services Unit of Sorbonne University.

### Oral glucose tolerance test (OGTT)

After 13 weeks of diet, overnight-fasted mice were treated with an oral gavage glucose load (2 g glucose per kg of body weight). Blood glucose was measured at time 0 just before oral glucose load and then 15, 30, 60 and 90 min after oral glucose load. Blood glucose was determined with a glucose meter (Accu Check, Roche, Switzerland) on blood samples collected from the tip of the tail vein. Plasma insulin concentration was determined using an ELISA kit (Mercodia, Uppsala, Sweden) according to the manufacturer’s instructions. HOMA-IR index was calculated according to the formula: fasting insulin (microU/L) x fasting glucose (nmol/L)/22.5 (Haffner et al. 1996).

### Insulin tolerance test (ITT)

After 14 weeks of diet, mice were fasted for six hours and blood glucose levels were determined before and at 15, 30, 60 and 90 min post an intraperitoneal injection of regular human insulin (Humulin®, Lilly; Indianapolis, IN - 0.75 U per kg of body weight).

### Adipocyte measurement

The mean adipocyte diameters were determined from subcutaneous adipose tissue after 24 weeks of HFD as previously described(Prat-Larquemin et al. 2004). Briefly, subcutaneous adipose tissue was rapidly washed with physiologic saline and then incubated with collagenase (1 mg/mL – Sigma-Aldrich, St. Quentin Fallavier, France) in phosphate buffer saline solution (pH 7.4) at 37°C for 20 minutes. An aliquot of floating mature adipocyte suspension was placed in a circular silicone ring (0.5 cm diameter) that was fixed to a silicon glass slide, to limit the dispersion of the adipocyte suspension, and then visualized under a light optic microscope connected to a camera and computer interface. Adipocyte diameters were measured with PERFECT IMAGE software (Numeris). Mean diameter was defined as the mean value for the distribution of adipocyte diameters of 150 cells.

### Intestinal paracellular permeability test

Intestinal paracellular permeability measurement in vivo was based on the intestinal permeability to 4 000 Da fluorescent dextran-FITC (Sigma-Aldrich). FITC-dextran was administered by gavage to 6-h-fasted-mice (600 mg per kg of body weight). 1h and 3h post-gavage, blood was collected from the tip of the tail vein (40µl) into EDTA-coated tubes and centrifuged (4°C, 2 000 g for 10 min). Plasma was diluted 1:10 (v/v) in phosphate buffered saline (PBS, pH 7.4) and the dextran-FITC concentration was determined using a fluorescence spectrophotometer (Fluostar; SLT, Crailsheim, Germany) at 485 nm excitation and 535 nm emission wavelengths. Standard curves were obtained by diluting dextran-FITC in non-treated plasma prepared in PBS (1:10 v/v).

### Fecal calorimetry

Twenty weeks after the beginning of the diet, one week of feces was collected per cage of 3 to 4 mice each. During the same time the food intake was monitored. The feces were dried overnight at 70°C and weighted. Total energy content of the feces was determined by bomb calorimetry (C200 bomb calorimeter, IKA compagny, Staufen, Germany) and results were expressed as kcal/day.

### Quantification of Fecal Lcn-2 by ELISA

Frozen fecal pellets were reconstituted in PBS containing 0.1% Tween 20 (100 mg/ml) and vortexed for 20 min. Fecal homogenates were then centrifuged for 10 min at 10,000 g and 4°C. Clear supernatants were collected and stored at −20°C until analysis. Lcn-2 levels were quantified in the supernatants using the Duoset murine Lcn-2 ELISA kit (R&D Systems, Minneapolis, MN).

### Quantification of Fecal albumin by ELISA

Feces were reconstituted in PBS (100mg/ml) and vortexed for 20 min. Fecal homogenates were then centrifuged for 10 min at 10,000 g and 4°C. Clear supernatants were collected and stored at −20°C until analysis. Albumin content in the feces was determined by ELISA following manufacturer’s instructions (Bethyl Laboratories, Montgomery, AL, USA).

### Quantification of Fecal SIgA by ELISA

Frozen fecal samples diluted 5-fold (w/v) in protease inhibitor cocktail containing PMSF (5mM), EDTA (1mM) and pepstatin (1 µg/mL - Sigma-Aldrich) were homogenized and centrifuged 10 min at 10,000 g at 4°C to collect supernatants. Flat bottom 96-well plates (Immulon II, VWR) coated with 100 µL/well of goat anti-mouse IgA (5 µg/mL Bic 0.1M, pH 9.6; Southern Biotech) were incubated with serial 3-fold dilutions of either fecal supernatant (100 µL) or standard IgA (100 µL - SouthernBiotech, Birmingham, USA) for 1h30 at 37°C. After washing, fixed antibodies were detected with horseradish peroxidase-conjugated goat anti-mouse IgA (100 µL / well - 1.5 µg/mL; Sigma-Aldrich) for 1h30 at 37°C and the reaction revealed with of 3,3’-5,5’-tetramethylbenzidine peroxidase substrate (100 µL/well - KPL, VWR, Fontenay-sous-Bois, France). Absorbencies were read at 450 nm.

### Fecal flagellin and LPS load quantification

Flagellin and Lipopolysaccharide (LPS) were quantified as previously described(Chassaing et al. 2014). We quantified flagellin and LPS using human embryonic kidney (HEK)-Blue-mouse (m)TLR5 and HEK-Blue-mTLR4 cells, respectively (Invivogen, San Diego, California, USA). We resuspended fecal material in PBS to a final concentration of 100 mg/mL and homogenized for 10 s using a Mini-Beadbeater-24 without the addition of beads to avoid bacteria disruption. We then centrifuged the samples at 8000 g for 2 min and serially diluted the resulting supernatant and applied to mammalian cells. Purified E coli flagellin and LPS (Sigma, St Louis, Missouri, USA) were used as positive controls for HEK-Blue-mTLR5 and HEK-Blue-mTLR4 cells, respectively. After 24 h of stimulation, we applied cell culture supernatant to QUANTI-Blue medium (Invivogen) and measured alkaline phosphatase activity at 620 nm after 30 min.

### Cytokine Secretion Assay

Single cell suspensions were prepared from mLNs, SILP and CLP.

After surgical removal of small intestine and colon, the SILP and the CLP single cell suspensions were obtained using the Lamina Propria Dissociation Kit (Miltenyi Biotec SAS, Paris, France) following the manufacturer’s instructions. Leucocytes enrichment was then performed through a 40/80% (w/v) Percoll density gradient (GE Healthcare) centrifuged for 15min at 1900g at RT. mLNs were surgically removed and then thoroughly smashed on a 70µm cell strainer on ice.

Single cell preparations were washed prior and resuspended in a complete media containing DMEM-Glutamax added with 8% fetal calf serum (FCS; PAA Laboratories, Linz, Austria), HEPES (10 mM), 2-mercaptoethanol (0.05 mM), and of penicillin and streptomycin (100 U/ml).

Single cell preparations were cultured in complete media (10^6^ cells /ml) for 72h in anti-CD3/anti-CD28 (5 μg/mL; BD Biosciences, San Jose, USA) pre-coated 96-flat-well plates (BD Biosciences). Supernatants were harvested and cytokine secretion assessed using BioPlex assay (Luminex MAGPIX Instrument, Bio-Rad, Marne-la-Coquette, France) according to manufacturer’s instructions.

### Flow Cytometry and Cell Sorting

Single cell preparations were pre-incubated with Fc-Block (eBioscience, Thermo Fisher Scientific, Les Ulis, France) for 20 min at 4°C. To differentiate live cells from dead cells, they were incubated 30 min at 4°C with fixable viability dye eFluor 506 (eBioscience). Cells were further stained for 30 min with antibodies to surface markers, the antibodies used are available in the Supplementary Material. For intracellular staining, cells were fixed and permeabilized with a commercially available fixation/permeabilization buffer (eBioscience). Intracellular staining was performed with PE-conjugated Foxp3 (clone FJK-16s) or with BV421-conjugated IFNg (clone XMG1.2) and APC-conjugated IL10 (clone JES516E3) and PE-cyanine7-conjugated IL17A (clone eBio17B7) or with v450-conjugated hBcl2 (clone Bcl-2/100).

Prior intracellular cytokine staining, cells were restimulated with PMA (50 ng/ml) and ionomycin (500 ng/ml) for 4 h in RPMI 1640 (Invitrogen) containing 8% FCS, HEPES (10 mM), 2-mercaptoethanol (0.05 mM), penicillin and streptomycin (100 U/ml).

Labelled cells were analyzed with a BD LSRFortessa flow cytometer (BD Biosciences) using both Diva or Flow-Jo software. Cell sorting experiments were performed on single cell preparations from mLNs of 8-weeks old mice before starting the HFD. After surface staining, CD103^+^ CD11b^+^ cDCs were Fluorescence-activated cell sorting (FACS)-sorted with the MoFlo Astrios EQ cell analyzer (Beckman Coulter).

### Analysis of RALDH activity by aldefluor staining

RALDH activity in individual cells was analyzed using the aldefluor staining kit (StemCell Technologies, Vancouver, BC, Canada). Briefly, 1.10^6^ cells were resuspended in the kit Assay Buffer containing activated aldefluor substrate (150 nM) and incubated for 30 min at 37°C in the presence or absence of the RALDH inhibitor DEAB (100 mM). Afterward cells were washed, placed on ice, stained for surface markers and analyzed through flow cytometry.

### 16S rRNA gene sequencing and analysis

Feces were collected at day 0 and 12 weeks after starting the diet and immediately frozen in liquid nitrogen and then stored at −80°C. Fecal DNA was extracted as previously described (Godon et al. 1997). The V3-V4 region of the 16S rRNA gene was amplified with the universal primers F343 (CTTTCCCTACACGACGCTCTTCCGATCTACGGRAGGCAGCAG) and R784 (GGAGTTCAGACGTGTGCTCTTCCGATCTTACCAGGGTATCTAATCCT), using 30 amplification cycles with an annealing temperature of 65 °C. The resulting PCR products were purified and sequenced at the GeT-PlaGe Genotoul INRA platform (Toulouse, France) using 506 Illumina MiSeq technology. The sequences were demultiplexed and quality filtered using the Quantitative Insights Into Microbial Ecology (QIIME, version 1.8.0) software package(Caporaso et al. 2010). We used QIIME default parameters for quality filtering (reads truncated at first low-quality base and excluded if: (1) there were more than three consecutive low quality base calls; (2) less than 75% of read length was consecutive high quality base calls; (3) at least one uncalled base was present; (4) more than 1.5 errors were present in the barcode; (5) any Phred qualities were below 20; or (6) the length was less than 75 bases). Sequences were assigned to OTUs using the UCLUST algorithm(Edgar 2010) with a 97% threshold of pairwise identity and without the creation of new clusters with sequences that did not match the reference sequences. OTUs were taxonomically classified using the Greengenes 13_8 reference database (McDonald et al. 2012). A single representative sequence for each OTU was aligned and a phylogenetic tree was built using FastTree (Price, Dehal, and Arkin 2009). The phylogenetic tree was used for computing the unweighted UniFrac distances between samples (Lozupone, Hamady, and Knight 2006; Lozupone and Knight 2005). Rarefied OTU table were used to compare abundances of OTUs across samples. Principal component analysis (PCA) plots were used to assess the variation between experimental group (beta diversity), alpha diversity curves were determined for all samples using the determination of the number of observed species, and OTU table was rarefied at various taxonomic levels using QIIME. LEfSE (LDA Effect Size) was used to investigate bacterial members that drive differences between groups (Segata et al. 2011). Unprocessed sequencing data are deposited in the European Nucleotide Archive under accession numbers XXXXX.

### Fecal Microbiota Transplantation

Feces from donor mice were diluted (30-50 mg - 1:10 w/vol) and homogenized in reduced sterile Ringer solution (VWR) containing L-Cysteine (0.5 g/L - Sigma-Aldrich) as reducing agent. This solution was immediately administered to germ-free recipients by oral gavage. Eight-weeks old germ-free mice were inoculated with donor fecal microbiota immediately after the opening of their sterile shipping container and once per week during the first three weeks of HFD. Recipient mice were then fed a HFD for 24 weeks. Recipients from WT donor are referred to as the FT-WT group, the recipients from DC^hBcl-2^ are referred to as FT-DC^hBcl-2^ (n = 12 mice per group).

### Short-chain fatty acid analysis in fecal and caecal samples

Samples were water extracted and proteins were precipitated with phosphotungstic acid. A volume of 0·1µl supernatant fraction was analyzed for SCFA on a gas–liquid chromatograph (Autosystem XL; Perkin Elmer, Saint-Quentin-en-Yvelines, France) equipped with a split-splitless injector, a flame-ionisation detector and a capillary column (15 m x 0·53 mm, 0·5µm) impregnated with SP 1000 (FSCAP Supelco, Saint-Quentin-Fallavier, France). Carrier gas (He) flow rate was 10 ml/min and inlet, column and detector temperatures were 175°C, 100°C and 280°C, respectively. 2-Ethylbutyrate was used as the internal standard (Lan et al. 2007). Samples were analysed in duplicate. Data were collected and peaks integrated using the Turbochrom v. 6 software (Perkin Elmer, Courtaboeuf, France).

### Microarray analysis

After FACS-cell sorting, cells were counted and resuspended in Trizol lysis reagent (Thermo Fisher Scientific), frozen in liquid nitrogen, and stored at −80°C. RNA extraction was performed using the RNeasy micro kit (Qiagen) following the manufacturer’s instructions. Quality and quantity of RNA extraction was performed using the Bioanalyzer 2100 RNA 6000 pico chip assay (Agilent). Total RNA (2.5 ng) was reverse transcribed following the Ovation Pico WTA System V2 (Nugen). cDNA hybridization was performed using the GeneChip® Mouse Gene 2.0 ST (Affymetrix) following the manufacturer’s instructions. Raw data (CEL files) were quality controlled, normalized and processed into signal intensities using the RMA algorithm with Affymetrix CDF file used for annotation. All subsequent analyses were based on the log (base 2) transformed data in Partek Genomics Suite: non-supervised analysis and Anova were used to detect eventual outlier samples and to identify differentially expressed genes. Statistical and Hierarchial clustering was performed using the TIGR Multiple Experiment Viewer (MeV 4.9.0) (Ai et al. 2003). Comparisons of the two groups were performed by a 2 two-tailed Student’s t-test. Features were considered significant when the p value was below 0.05 after Benjamini-Hochberg for false discovery rate (FDR) correction. The resulted DEGs were mapped for Gene Ontology (GO) and KEGG/BioCarta pathway analysis using ClueGO (Bindea et al. 2009) (version 2.5.5) a Cytoscape (Shannon et al. 2003) (version 3.7.0) plug-in facilitating the biological interpretation and visualization of functionally grouped GO terms in the form of networks and charts. A two-sided (enrichment/depletion) hyper-geometric distribution test with a p-value significance level of ≤ 0.05 corrected by Bonferroni were applied, together with the Kappa-statistic score threshold at 0.3 and GO levels set between 4 to 6. Datasets were derived from 8 to 12 samples per genotype. 2 to 3 samples were pooled for each genotype (corresponding to either group 1, 2, 3 or 4 in the figure) to obtain equivalent amount of material for further processing. Each group of pooled samples correspond to Gp1 to Gp4 (Figure 3D). All original microarray data were deposited in the NCBI’s Gene Expression Omnibus database (GEO XXXXXXXXX).

### Statistical analysis

Data are expressed as medians for dot plot or mean and SD for bar plots. Data were analyzed using GraphPad Prism version 8.00 for windows (GraphPad Software, San Diego, CA, USA). Mann-Whitney test and Kruskal-Wallis test or two-ways ANOVA for multiple comparisons were performed. Results were considered statistically significant when *p* < 0.05. Stars indicate significant differences (*: *p* < 0.05; **: *p* < 0.01; ***: *p* < 0.001) between two groups according to statistical analysis performed.

## Supporting Information

### Indirect calorimetry

A subset of WT and DC^hBcl-2^ mice HFD-fed for 20 weeks was used for indirect calorimetry measurement (n = 6 per group). These mice were housed individually in metabolic chambers (Phenomaster, TSE Systems GmbH, Bad Homburg, Germany). After 3 days of habituation, the measurement of food intake, drink intake, locomotor activity, O2 consumption, CO2 production and energy expenditure were monitored for 5 days.

### Transit time

Carmine red was given by gavage to 6-h-fasted mice (10 mg/ml of water, 10 µl per g of body weight). The intestinal transit time was recorded as the time from gavage to the first appearance of the dye in the feces (minutes).

### Cell surface staining

Surface staining was performed using the following antibodies (BD Biosciences): FITC-CD45 (clone 1D3), Alexa Fluor-anti-B220 (clone RA3-6B/2), v450-anti-CD19 (clone1D3), Alexa-Fluor 700-anti-MHC Class II (I-A/I-E) (clone M5/114.15.2), PE-Cy7-anti-CD11c (clone HL3), Allophycocyanin (APC)-anti-CD64 (clone X54–5/7.1), APC-Cy7-anti-CD11b (clone M1/70), PE-anti-CD103 (clone M290), PerCP-Cy5.5-anti-CD3 (clone 17A2), PerCP-eFluor710-anti-CD3 (clone 17A2), BV-711-anti-CD4 (RM4-5), PE-CF594-anti-CD8a (clone 53-6.7), PE-anti-IgA, (SouthernBiotech).

## Acknowledgements

This study was supported by the Institut National de la Santé et de la Recherche Médicale (INSERM), Sorbonne Université (SU), the Fondation de France (00029519), and the Institute of Cardiometabolism and Nutrition (IHU-ICAN, ANR-10-IAHU-05). E.L. was supported by the Fondation Lefoulon Delalande/Institut de France and the Region Ile-de-France CORDDIM.

We are grateful to the PreclinICAN and CytoICAN platforms from IHU-ICAN, the “plate-forme de Génomique, Institut Cochin, Inserm 1016-CNRS 8104-Paris Descartes” and the animal facility of “Centre d’expérimentation fonctionnelle, Equipe du 105B, La Pitié-Salpêtrière, Paris” for excellent technical support. We also thank François Déjardin and Julien Verdier for fruitful discussions and careful reading of the manuscript.

## Competing Interests

The authors declare no conflict of interest.

### Author Contributions

Designed the experiments: E.L., T.L.R., A.G., A.L., N.V., M.G. and P.L. Performed the experiments: E.L., T.L.R., A.G., A.L., J.B.H., C.P., M.F., F.I., S.B., M.R., E.M., N.K., P.G., B.C. and P.L. Performed the analysis: E.L., T.L.R., A.G., A.L., J.B.H., C.P., M.P., N.K., and B.C. Provided resources: E.L., M.G., P.L., Drafted the manuscript: E.L.,T.L.R., M.P. and B.C. Revised the manuscript: E.L., T.L.R., M.F., F.I., T.H., E.G., N.K., P.G., N.V., B.C. and P.L.

## References

Ai, Saeed, Sharov V, White J, Li J, Liang W, Bhagabati N, Braisted J, et al. 2003. “TM4: A Free, Open-Source System for Microarray Data Management and Analysis.” BioTechniques. Biotechniques. February 2003. https://doi.org/10.2144/03342mt01.

Arpaia, Nicholas, Clarissa Campbell, Xiying Fan, Stanislav Dikiy, Joris van der Veeken, Paul deRoos, Hui Liu, et al. 2013. “Metabolites Produced by Commensal Bacteria Promote Peripheral Regulatory T-Cell Generation.” Nature 504 (7480): 451–55. https://doi.org/10.1038/nature12726.

Bai, Aiping, Nonghua Lu, Yuan Guo, Zhanju Liu, Jiang Chen, and Zhikang Peng. 2009. “All-Trans Retinoic Acid down-Regulates Inflammatory Responses by Shifting the Treg/Th17 Profile in Human Ulcerative and Murine Colitis.” Journal of Leukocyte Biology 86 (4): 959–69. https://doi.org/10.1189/jlb.0109006.

Bekiaris, Vasileios, Emma K. Persson, and William W. Agace. 2014. “Intestinal Dendritic Cells in the Regulation of Mucosal Immunity.” Immunological Reviews 260 (1): 86–101. https://doi.org/10.1111/imr.12194.

Belkaid, Yasmine, and Timothy Hand. 2014. “Role of the Microbiota in Immunity and Inflammation.” Cell 157 (1): 121–41. https://doi.org/10.1016/j.cell.2014.03.011.

Benjamini, Yoav, and Yosef Hochberg. 1995. “Controlling the False Discovery Rate: A Practical and Powerful Approach to Multiple Testing.” Journal of the Royal Statistical Society. Series B (Methodological) 57 (1): 289–300.

Bindea, Gabriela, Bernhard Mlecnik, Hubert Hackl, Pornpimol Charoentong, Marie Tosolini, Amos Kirilovsky, Wolf-Herman Fridman, Franck Pagès, Zlatko Trajanoski, and Jérôme Galon. 2009. “ClueGO: A Cytoscape Plug-in to Decipher Functionally Grouped Gene Ontology and Pathway Annotation Networks.” Bioinformatics 25 (8): 1091–93. https://doi.org/10.1093/bioinformatics/btp101.

Brown, Eric M., Manish Sadarangani, and B. Brett Finlay. 2013. “The Role of the Immune System in Governing Host-Microbe Interactions in the Intestine.” Nature Immunology 14 (7): 660–67. https://doi.org/10.1038/ni.2611.

Cani, Patrice D., Jacques Amar, Miguel Angel Iglesias, Marjorie Poggi, Claude Knauf, Delphine Bastelica, Audrey M. Neyrinck, et al. 2007a. “Metabolic Endotoxemia Initiates Obesity and Insulin Resistance.” Diabetes 56 (7): 1761–72. https://doi.org/10.2337/db06-1491.

Cani, Patrice D. 2007b. “Metabolic Endotoxemia Initiates Obesity and Insulin Resistance.” Diabetes 56 (7): 1761–72. https://doi.org/10.2337/db06-1491.

Cani, Patrice D., Rodrigo Bibiloni, Claude Knauf, Aurélie Waget, Audrey M. Neyrinck, Nathalie M. Delzenne, and Rémy Burcelin. 2008. “Changes in Gut Microbiota Control Metabolic Endotoxemia- Induced Inflammation in High-Fat Diet-Induced Obesity and Diabetes in Mice.” Diabetes 57 (6): 1470–81. https://doi.org/10.2337/db07-1403.

Caporaso, J Gregory, Justin Kuczynski, Jesse Stombaugh, Kyle Bittinger, Frederic D Bushman, Elizabeth K Costello, Noah Fierer, et al. 2010. “QIIME Allows Analysis of High-Throughput Community Sequencing Data.” Nature Methods 7 (5): 335–36. https://doi.org/10.1038/nmeth.f.303.

Cassani, Barbara, Eduardo J. Villablanca, Jaime De Calisto, Sen Wang, and J. Rodrigo Mora. 2012. “Vitamin A and Immune Regulation: Role of Retinoic Acid in Gut-Associated Dendritic Cell Education, Immune Protection and Tolerance.” Molecular Aspects of Medicine 33 (1): 63–76. https://doi.org/10.1016/j.mam.2011.11.001.

Chang, Sun-Young, Hyun-Jeong Ko, and Mi-Na Kweon. 2014. “Mucosal Dendritic Cells Shape Mucosal Immunity.” Experimental & Molecular Medicine 46 (March): e84. https://doi.org/10.1038/emm.2014.16.

Chassaing, Benoit, Omry Koren, Frederic A. Carvalho, Ruth E. Ley, and Andrew T. Gewirtz. 2014. “AIEC Pathobiont Instigates Chronic Colitis in Susceptible Hosts by Altering Microbiota Composition.” Gut 63 (7): 1069–80. https://doi.org/10.1136/gutjnl-2013-304909.

Chassaing, Benoit, Gayathri Srinivasan, Maria A. Delgado, Andrew N. Young, Andrew T. Gewirtz, and Matam Vijay-Kumar. 2012. “Fecal Lipocalin 2, a Sensitive and Broadly Dynamic Non-Invasive Biomarker for Intestinal Inflammation.” PloS One 7 (9): e44328. https://doi.org/10.1371/journal.pone.0044328.

Coombes, Janine L., and Fiona Powrie. 2008. “Dendritic Cells in Intestinal Immune Regulation.” Nature Reviews. Immunology 8 (6): 435–46. https://doi.org/10.1038/nri2335.

Coombes, Janine L., Karima R. R. Siddiqui, Carolina V. Arancibia-Cárcamo, Jason Hall, Cheng- Ming Sun, Yasmine Belkaid, and Fiona Powrie. 2007. “A Functionally Specialized Population of Mucosal CD103+ DCs Induces Foxp3+ Regulatory T Cells via a TGF-β– and Retinoic Acid– Dependent Mechanism.” Journal of Experimental Medicine 204 (8): 1757–64. https://doi.org/10.1084/jem.20070590.

De Vadder, Filipe, Petia Kovatcheva-Datchary, Daisy Goncalves, Jennifer Vinera, Carine Zitoun, Adeline Duchampt, Fredrik Bäckhed, and Gilles Mithieux. 2014. “Microbiota-Generated Metabolites Promote Metabolic Benefits via Gut-Brain Neural Circuits.” Cell 156 (1–2): 84–96. https://doi.org/10.1016/j.cell.2013.12.016.

Ding, Shengli, Michael M. Chi, Brooks P. Scull, Rachael Rigby, Nicole M. J. Schwerbrock, Scott Magness, Christian Jobin, and Pauline K. Lund. 2010. “High-Fat Diet: Bacteria Interactions Promote Intestinal Inflammation Which Precedes and Correlates with Obesity and Insulin Resistance in Mouse.” PLOS ONE 5 (8): e12191. https://doi.org/10.1371/journal.pone.0012191.

Edgar, Robert C. 2010. “Search and Clustering Orders of Magnitude Faster than BLAST.” Bioinformatics (Oxford, England) 26 (19): 2460–61. https://doi.org/10.1093/bioinformatics/btq461.

Fernández-Ruiz, Irene. 2016. “Immune System and Cardiovascular Disease.” Nature Reviews Cardiology 13 (9): 503–503. https://doi.org/10.1038/nrcardio.2016.127.

Gao, Zhanguo, Jun Yin, Jin Zhang, Robert E. Ward, Roy J. Martin, Michael Lefevre, William T. Cefalu, and Jianping Ye. 2009. “Butyrate Improves Insulin Sensitivity and Increases Energy Expenditure in Mice.” Diabetes 58 (7): 1509–17. https://doi.org/10.2337/db08-1637.

Garidou, Lucile, Céline Pomié, Pascale Klopp, Aurélie Waget, Julie Charpentier, Meryem Aloulou, Anaïs Giry, et al. 2015. “The Gut Microbiota Regulates Intestinal CD4 T Cells Expressing RORγt and Controls Metabolic Disease.” Cell Metabolism 22 (1): 100–112. https://doi.org/10.1016/j.cmet.2015.06.001.

Gautier, Emmanuel L., Thierry Huby, Flora Saint-Charles, Betty Ouzilleau, M. John Chapman, and Philippe Lesnik. 2008. “Enhanced Dendritic Cell Survival Attenuates Lipopolysaccharide-Induced Immunosuppression and Increases Resistance to Lethal Endotoxic Shock.” The Journal of Immunology 180 (10): 6941. https://doi.org/10.4049/jimmunol.180.10.6941.

Gautier Emmanuel L., Huby Thierry, Saint-Charles Flora, Ouzilleau Betty, Pirault John, Deswaerte Virginie, Ginhoux Florent, et al. 2009. “Conventional Dendritic Cells at the Crossroads Between Immunity and Cholesterol Homeostasis in Atherosclerosis.” Circulation 119 (17): 2367–75. https://doi.org/10.1161/CIRCULATIONAHA.108.807537.

Godon, J J, E Zumstein, P Dabert, F Habouzit, and R Moletta. 1997. “Molecular Microbial Diversity of an Anaerobic Digestor as Determined by Small-Subunit RDNA Sequence Analysis.” Applied and Environmental Microbiology 63 (7): 2802–13.

Granucci, Francesca, Caterina Vizzardelli, Norman Pavelka, Sonia Feau, Maria Persico, Ettore Virzi, Maria Rescigno, Giorgio Moro, and Paola Ricciardi-Castagnoli. 2001. “Inducible IL-2 Production by Dendritic Cells Revealed by Global Gene Expression Analysis.” Nature Immunology 2 (9): 882–88. https://doi.org/10.1038/ni0901-882.

Haffner, S. M., C. Gonzalez, H. Miettinen, E. Kennedy, and M. P. Stern. 1996. “A Prospective Analysis of the HOMA Model: The Mexico City Diabetes Study.” Diabetes Care 19 (10): 1138–41. https://doi.org/10.2337/diacare.19.10.1138.

Hong, Chun-Pyo, Areum Park, Bo-Gie Yang, Chang Ho Yun, Min-Jung Kwak, Gil-Woo Lee, Jung-Hwan Kim, et al. 2017. “Gut-Specific Delivery of T-Helper 17 Cells Reduces Obesity and Insulin Resistance in Mice.” Gastroenterology 152 (8): 1998–2010. https://doi.org/10.1053/j.gastro.2017.02.016.

Hou, Wu-Shiun, and Luk Van Parijs. 2004. “A Bcl-2-Dependent Molecular Timer Regulates the Lifespan and Immunogenicity of Dendritic Cells.” Nature Immunology 5 (6): 583–89. https://doi.org/10.1038/ni1071.

Iwata, Makoto, Asami Hirakiyama, Yuko Eshima, Hiroyuki Kagechika, Chieko Kato, and Si-Young Song. 2004. “Retinoic Acid Imprints Gut-Homing Specificity on T Cells.” Immunity 21 (4): 527–38. https://doi.org/10.1016/j.immuni.2004.08.011.

Kaisar, Maria M. M., Leonard R. Pelgrom, Alwin J. van der Ham, Maria Yazdanbakhsh, and Bart Everts. 2017. “Butyrate Conditions Human Dendritic Cells to Prime Type 1 Regulatory T Cells via Both Histone Deacetylase Inhibition and G Protein-Coupled Receptor 109A Signaling.” Frontiers in Immunology 8 (October). https://doi.org/10.3389/fimmu.2017.01429.

Kawano, Yoshinaga, Jun Nakae, Nobuyuki Watanabe, Tetsuhiro Kikuchi, Sanshiro Tateya, Yoshikazu Tamori, Mari Kaneko, Takaya Abe, Masafumi Onodera, and Hiroshi Itoh. 2016. “Colonic Pro-Inflammatory Macrophages Cause Insulin Resistance in an Intestinal Ccl2/Ccr2-Dependent Manner.” Cell Metabolism 24 (2): 295–310. https://doi.org/10.1016/j.cmet.2016.07.009.

Laffont, Sophie, Karima R. R. Siddiqui, and Fiona Powrie. 2010. “Intestinal Inflammation Abrogates the Tolerogenic Properties of MLN CD103+ Dendritic Cells.” European Journal of Immunology 40 (7): 1877–83. https://doi.org/10.1002/eji.200939957.

Lan, Annaïg, Aurélia Bruneau, Catherine Philippe, Violaine Rochet, Annette Rouault, Christophe Hervé, Nathalie Roland, Sylvie Rabot, and Gwénaël Jan. 2007. “Survival and metabolic activity of selected strains of Propionibacterium freudenreichii in the gastrointestinal tract of human microbiota-associated rats.” British Journal of Nutrition 97 (4): 714–24. https://doi.org/10.1017/S0007114507433001.

Le Roy, Tiphaine, Marta Llopis, Patricia Lepage, Aurélia Bruneau, Sylvie Rabot, Claudia Bevilacqua, Patrice Martin, et al. 2013. “Intestinal Microbiota Determines Development of Non-Alcoholic Fatty Liver Disease in Mice.” Gut 62 (12): 1787–94. https://doi.org/10.1136/gutjnl-2012-303816.

Li, Meng, Betty C. A. M. van Esch, Gerry T. M. Wagenaar, Johan Garssen, Gert Folkerts, and Paul A. J. Henricks. 2018. “Pro- and Anti-Inflammatory Effects of Short Chain Fatty Acids on Immune and Endothelial Cells.” European Journal of Pharmacology 831 (July): 52–59. https://doi.org/10.1016/j.ejphar.2018.05.003.

Lozupone, Catherine, Micah Hamady, and Rob Knight. 2006. “UniFrac – An Online Tool for Comparing Microbial Community Diversity in a Phylogenetic Context.” BMC Bioinformatics 7 (1): 371. https://doi.org/10.1186/1471-2105-7-371.

Lozupone, Catherine, and Rob Knight. 2005. “UniFrac: A New Phylogenetic Method for Comparing Microbial Communities.” Applied and Environmental Microbiology 71 (12): 8228–35. https://doi.org/10.1128/AEM.71.12.8228-8235.2005.

Luck, Helen, Sue Tsai, Jason Chung, Xavier Clemente-Casares, Magar Ghazarian, Xavier S. Revelo, Helena Lei, et al. 2015. “Regulation of Obesity-Related Insulin Resistance with Gut Anti-Inflammatory Agents.” Cell Metabolism 21 (4): 527–42. https://doi.org/10.1016/j.cmet.2015.03.001.

Magnusson, M. K., S. F. Brynjólfsson, A. Dige, H. Uronen-Hansson, L. G. Börjesson, J. L. Bengtsson, S. Gudjonsson, et al. 2016. “Macrophage and Dendritic Cell Subsets in IBD: ALDH+ Cells Are Reduced in Colon Tissue of Patients with Ulcerative Colitis Regardless of Inflammation.” Mucosal Immunology 9 (1): 171–82. https://doi.org/10.1038/mi.2015.48.

Mahnke, Karsten, Edgar Schmitt, Laura Bonifaz, Alexander H. Enk, and Helmut Jonuleit. 2002. “Immature, but Not Inactive: The Tolerogenic Function of Immature Dendritic Cells.” Immunology and Cell Biology 80 (5): 477–83. https://doi.org/10.1046/j.1440-1711.2002.01115.x.

Mantis, N. J., N. Rol, and B. Corthésy. 2011. “Secretory IgA’s Complex Roles in Immunity and Mucosal Homeostasis in the Gut.” Mucosal Immunology 4 (6): 603–11. https://doi.org/10.1038/mi.2011.41.

McDonald, Daniel, Morgan N Price, Julia Goodrich, Eric P Nawrocki, Todd Z DeSantis, Alexander Probst, Gary L Andersen, Rob Knight, and Philip Hugenholtz. 2012. “An Improved Greengenes Taxonomy with Explicit Ranks for Ecological and Evolutionary Analyses of Bacteria and Archaea.” The ISME Journal 6 (3): 610–18. https://doi.org/10.1038/ismej.2011.139.

Merad, Miriam, Priyanka Sathe, Julie Helft, Jennifer Miller, and Arthur Mortha. 2013. “The Dendritic Cell Lineage: Ontogeny and Function of Dendritic Cells and Their Subsets in the Steady State and the Inflamed Setting.” Annual Review of Immunology 31: 563–604. https://doi.org/10.1146/annurev-immunol-020711-074950.

Mora, J. Rodrigo, Makoto Iwata, Bertus Eksteen, Si-Young Song, Tobias Junt, Balimkiz Senman, Kevin L. Otipoby, et al. 2006. “Generation of Gut-Homing IgA-Secreting B Cells by Intestinal Dendritic Cells.” Science (New York, N.Y.) 314 (5802): 1157–60. https://doi.org/10.1126/science.1132742.

Mucida, Daniel, Yunji Park, Gisen Kim, Olga Turovskaya, Iain Scott, Mitchell Kronenberg, and Hilde Cheroutre. 2007. “Reciprocal TH17 and Regulatory T Cell Differentiation Mediated by Retinoic Acid.” Science (New York, N.Y.) 317 (5835): 256–60. https://doi.org/10.1126/science.1145697.

Natividad, Jane M., Bruno Lamas, Hang Phuong Pham, Marie-Laure Michel, Dominique Rainteau, Chantal Bridonneau, Gregory da Costa, et al. 2018. “Bilophila Wadsworthia Aggravates High Fat Diet Induced Metabolic Dysfunctions in Mice.” Nature Communications 9 (1): 1–15. https://doi.org/10.1038/s41467-018-05249-7.

Nopora, Adam, and Thomas Brocker. 2002. “Bcl-2 Controls Dendritic Cell Longevity In Vivo.” The Journal of Immunology 169 (6): 3006–14. https://doi.org/10.4049/jimmunol.169.6.3006.

O’Neill, S., and L. O’Driscoll. 2015. “Metabolic Syndrome: A Closer Look at the Growing Epidemic and Its Associated Pathologies.” Obesity Reviews: An Official Journal of the International Association for the Study of Obesity 16 (1): 1–12. https://doi.org/10.1111/obr.12229.

Pabst, O., and A. M. Mowat. 2012. “Oral Tolerance to Food Protein.” Mucosal Immunology 5 (3): 232–39. https://doi.org/10.1038/mi.2012.4.

Parada Venegas, Daniela, Marjorie K. De la Fuente, Glauben Landskron, María Julieta González, Rodrigo Quera, Gerard Dijkstra, Hermie J. M. Harmsen, Klaas Nico Faber, and Marcela A. Hermoso. 2019. “Short Chain Fatty Acids (SCFAs)-Mediated Gut Epithelial and Immune Regulation and Its Relevance for Inflammatory Bowel Diseases.” Frontiers in Immunology 10 (March). https://doi.org/10.3389/fimmu.2019.00277.

Persson, Emma K., Heli Uronen-Hansson, Monika Semmrich, Aymeric Rivollier, Karin Hägerbrand, Jan Marsal, Sigurdur Gudjonsson, et al. 2013. “IRF4 Transcription-Factor-Dependent CD103+CD11b+ Dendritic Cells Drive Mucosal T Helper 17 Cell Differentiation.” Immunity 38 (5): 958–69. https://doi.org/10.1016/j.immuni.2013.03.009.

Petersen, Charisse, Rickesha Bell, Kendra A. Klag, Soh-Hyun Lee, Raymond Soto, Arevik Ghazaryan, Kaitlin Buhrke, et al. 2019. “T Cell–Mediated Regulation of the Microbiota Protects against Obesity.” Science 365 (6451). https://doi.org/10.1126/science.aat9351.

Prat-Larquemin, Lydie, Jean-Michel Oppert, Karine Clément, Isabelle Hainault, Arnaud Basdevant, Bernard Guy-Grand, and Annie Quignard-Boulangé. 2004. “Adipose Angiotensinogen Secretion, Blood Pressure, and AGT M235T Polymorphism in Obese Patients.” Obesity Research 12 (3): 556–61. https://doi.org/10.1038/oby.2004.63.

Price, Morgan N., Paramvir S. Dehal, and Adam P. Arkin. 2009. “FastTree: Computing Large Minimum Evolution Trees with Profiles Instead of a Distance Matrix.” Molecular Biology and Evolution 26 (7): 1641–50. https://doi.org/10.1093/molbev/msp077.

Qiang, Y., J. Xu, C. Yan, H. Jin, T. Xiao, N. Yan, L. Zhou, et al. 2017. “Butyrate and Retinoic Acid Imprint Mucosal-like Dendritic Cell Development Synergistically from Bone Marrow Cells.” Clinical and Experimental Immunology 189 (3): 290–97. https://doi.org/10.1111/cei.12990.

Saklayen, Mohammad G. 2018. “The Global Epidemic of the Metabolic Syndrome.” Current Hypertension Reports 20 (2). https://doi.org/10.1007/s11906-018-0812-z.

Segata, Nicola, Jacques Izard, Levi Waldron, Dirk Gevers, Larisa Miropolsky, Wendy S Garrett, and Curtis Huttenhower. 2011. “Metagenomic Biomarker Discovery and Explanation.” Genome Biology 12 (6): R60–R60. https://doi.org/10.1186/gb-2011-12-6-r60.

Shannon, Paul, Andrew Markiel, Owen Ozier, Nitin S. Baliga, Jonathan T. Wang, Daniel Ramage, Nada Amin, Benno Schwikowski, and Trey Ideker. 2003. “Cytoscape: A Software Environment for Integrated Models of Biomolecular Interaction Networks.” Genome Research 13 (11): 2498–2504. https://doi.org/10.1101/gr.1239303.

Singh, Nagendra, Ashish Gurav, Sathish Sivaprakasam, Evan Brady, Ravi Padia, Huidong Shi, Muthusamy Thangaraju, et al. 2014. “Activation of Gpr109a, Receptor for Niacin and the Commensal Metabolite Butyrate, Suppresses Colonic Inflammation and Carcinogenesis.” Immunity 40 (1): 128–39. https://doi.org/10.1016/j.immuni.2013.12.007.

Sun, Cheng-Ming, Jason A. Hall, Rebecca B. Blank, Nicolas Bouladoux, Mohamed Oukka, J. Rodrigo Mora, and Yasmine Belkaid. 2007. “Small Intestine Lamina Propria Dendritic Cells Promote de Novo Generation of Foxp3 T Reg Cells via Retinoic Acid.” The Journal of Experimental Medicine 204 (8): 1775–85. https://doi.org/10.1084/jem.20070602.

Tisch, Roland. 2010. “Immunogenic Versus Tolerogenic Dendritic Cells: A Matter of Maturation.” International Reviews of Immunology 29 (2): 111–18. https://doi.org/10.3109/08830181003602515.

Wang, Li, Limeng Zhu, and Song Qin. 2019. “Gut Microbiota Modulation on Intestinal Mucosal Adaptive Immunity.” Journal of Immunology Research 2019 (October): 1–10. https://doi.org/10.1155/2019/4735040.

Wang, Lirui, Cristina Llorente, Phillipp Hartmann, An-Ming Yang, Peng Chen, and Bernd Schnabl. 2015. “Methods to Determine Intestinal Permeability and Bacterial Translocation during Liver Disease.” Journal of Immunological Methods 421 (June): 44–53. https://doi.org/10.1016/j.jim.2014.12.015.

Wang, Xiaoting, Naruhisa Ota, Paolo Manzanillo, Lance Kates, Jose Zavala-Solorio, Celine Eidenschenk, Juan Zhang, et al. 2014. “Interleukin-22 Alleviates Metabolic Disorders and Restores Mucosal Immunity in Diabetes.” Nature 514 (7521): 237–41. https://doi.org/10.1038/nature13564.

Winer, Daniel A., Helen Luck, Sue Tsai, and Shawn Winer. 2016. “The Intestinal Immune System in Obesity and Insulin Resistance.” Cell Metabolism 23 (3): 413–26. https://doi.org/10.1016/j.cmet.2016.01.003.

Zhao, Qing, and Charles O. Elson. 2018. “Adaptive Immune Education by Gut Microbiota Antigens.” Immunology 154 (1): 28–37. https://doi.org/10.1111/imm.12896.

Zlotnikov-Klionsky, Yael, Bar Nathansohn-Levi, Elias Shezen, Chava Rosen, Sivan Kagan, Liat Bar- On, Steffen Jung, et al. 2015. “Perforin-Positive Dendritic Cells Exhibit an Immuno-Regulatory Role in Metabolic Syndrome and Autoimmunity.” Immunity 43 (4): 776–87. https://doi.org/10.1016/j.immuni.2015.08.015.

