## Supplementary material for "Tolerogenic Dendritic Cells Shape a Transmissible Gut Microbiota that Protects from Metabolic Diseases": Figure Supplement

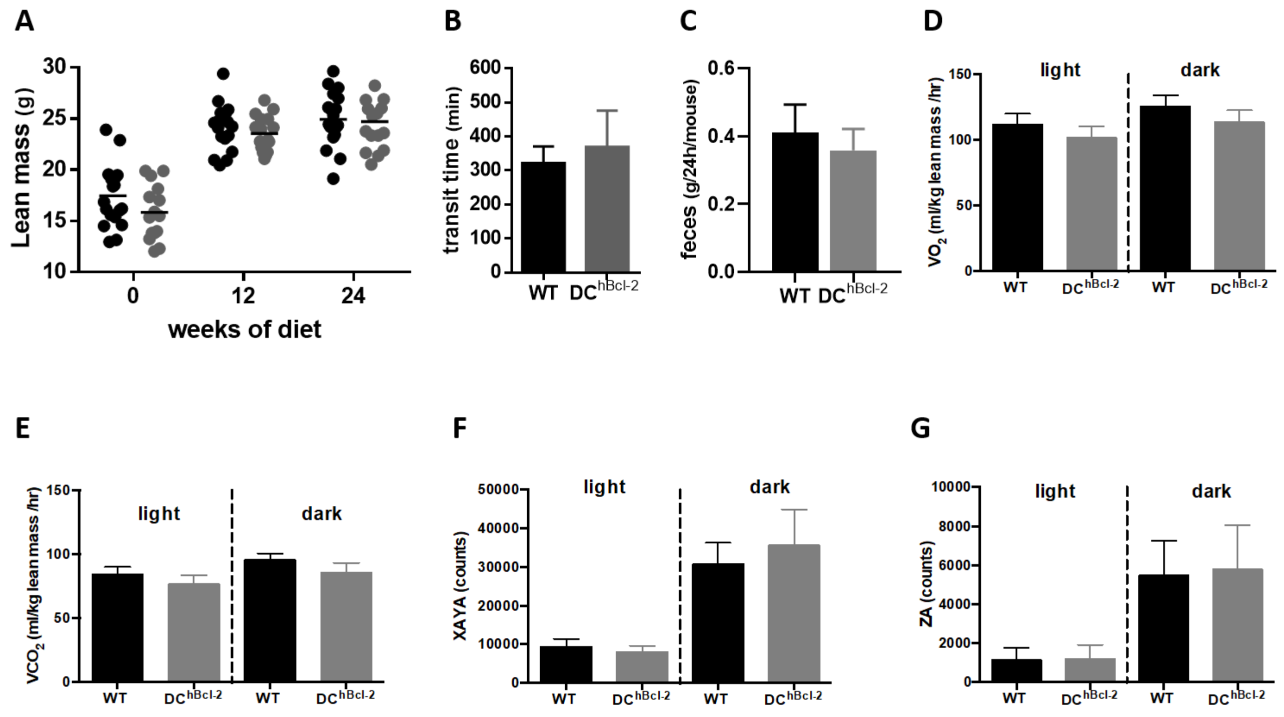


**figure supplement 1** (A) Lean mass monitoring at 0, 12 and 24 weeks after starting the HFD. (B) Transit time of mice after 24 weeks of HFD (N=7 to 9 mice per group). (C) Mean of feces production monitored for one week after 12-weeks of HFD. Volume of $O_2$ consumption (D) $CO_2$ produced (E) and ambulatory movements (F) and (G) of individually-housed mice in metabolic cages monitored for 5 days after 12-weeks of HFD (N=6 mice per group).

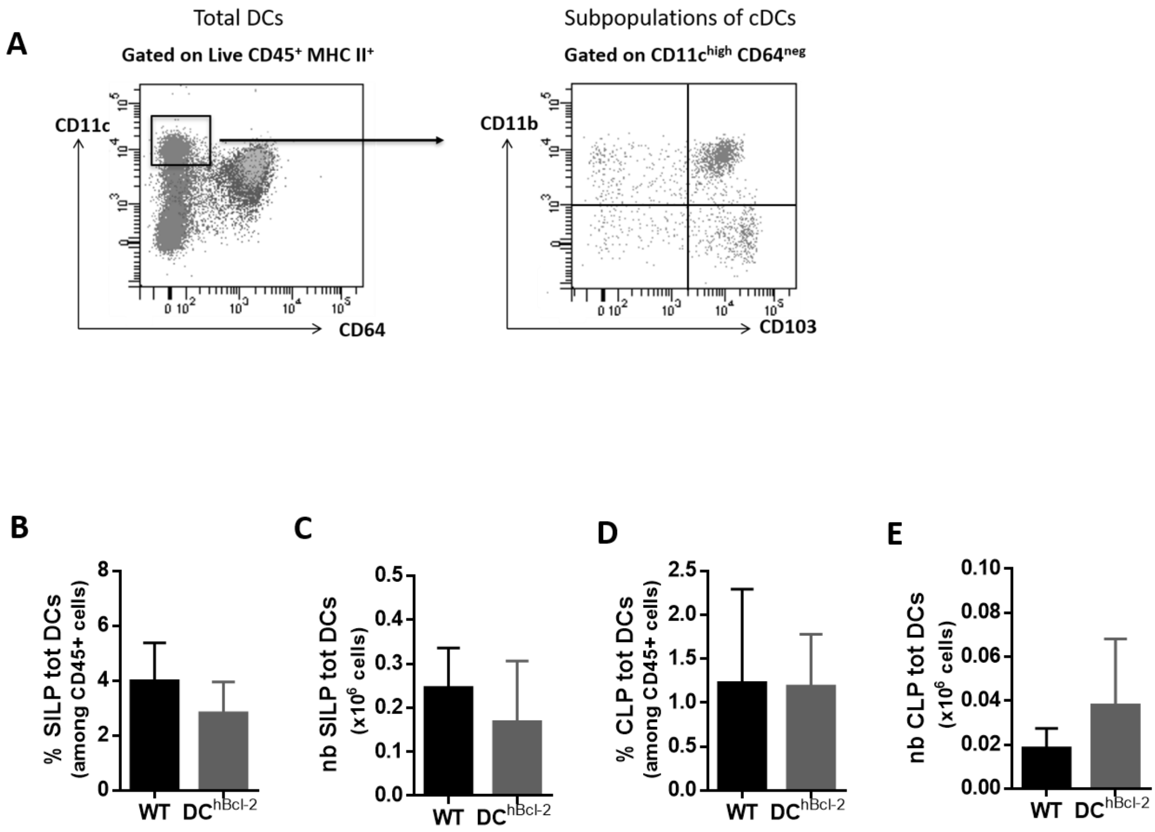


**figure supplement 2** (A) Gating strategy to target conventional DCs (cDCs) subpopulations in the intestine. Proportions of total dendritic cells (totDCs) after surface staining of cells in the SILP (C) or in the CLP (E) after 24 weeks of HFD (N=11 to 13 mice per group). Total numbers of total dendritic cells (totDCs) after surface staining of cells in the SILP (D) or in the CLP (F) after 24 weeks of HFD (N=11 to 13 mice per group).

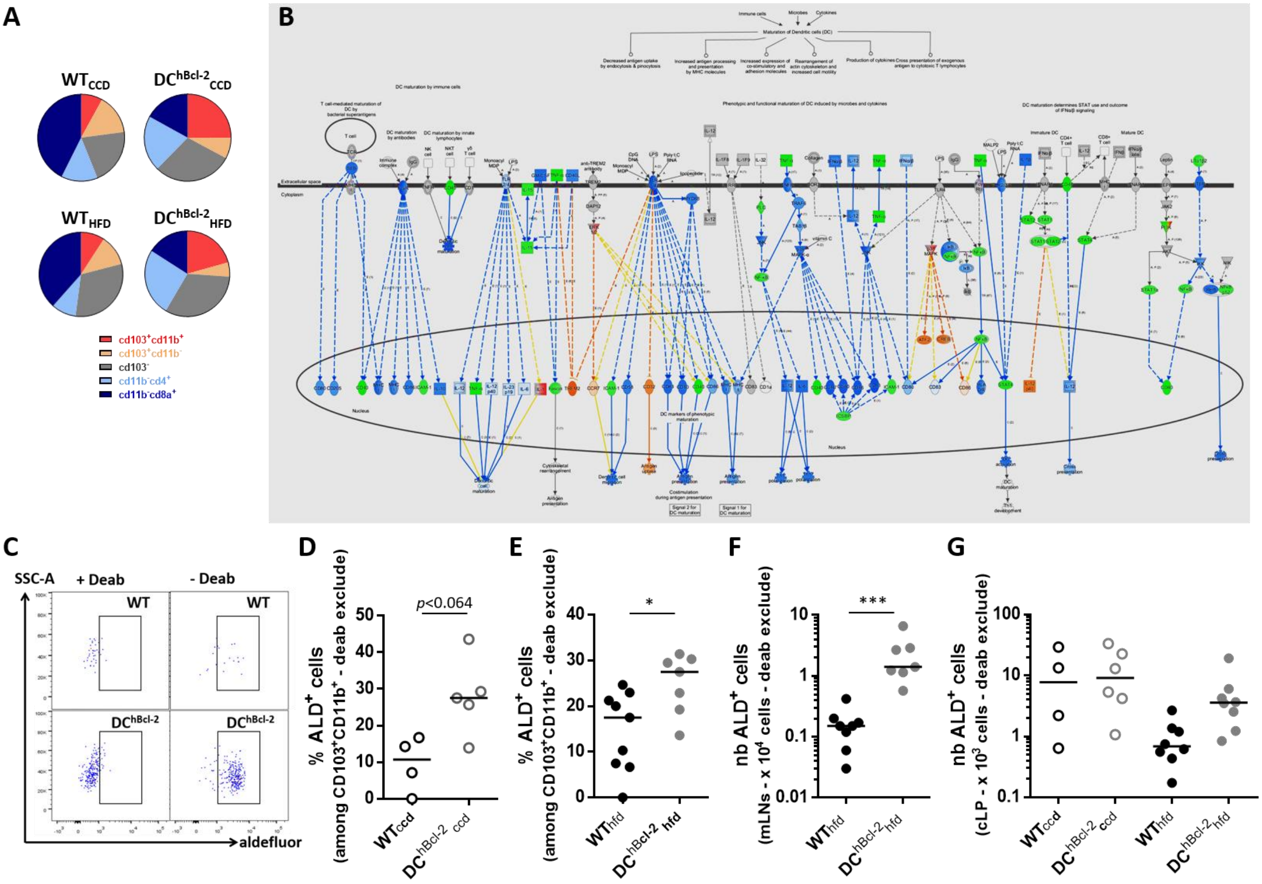


**figure supplement 3** (A) Circle graphs representing the mean proportions of conventional DCs subpopulations in the mLNs after 24 weeks of CCD or HFD. (B) Network diagram of DC maturation genes differentially expressed in DC$^{hBcl-2}$ mice relatively to WT mice, with corresponding predictive signaling pathways impacted using Ingenuity Pathway Analysis (IPA, Qiagen, Courtabœuf, France). The red or green colors indicate the degree of respective up-regulation or down-regulation in gene expression compared with housekeeping gene expression. The orange and the blue colors indicate the respective predictive degree of up-regulation or down-regulation in gene expression involved in these pathways. (C) Identification by FACS of the RALDH activity, i.e. aldefluor$^+$ (ALD$^+$) cells with or without the DEAB reagent (specific inhibitor for RALDH) in the CD103$^+$ CD11b$^+$ cDC subpopulations from the mLNs after 24 weeks of diet. Proportions of ALD$^+$ cells among CD103$^+$ CD11b$^+$ cDCs in the mLNs of mice after 24 weeks of CCD (D) or HFD (E). (F) Total numbers of CD103$^+$ CD11b$^+$ cDCs ALD$^+$ cells in the mLNs of mice after 24 weeks of HFD. (G) Total numbers of ALD$^+$ DCs in the CLP of mice after 24 weeks of diet.

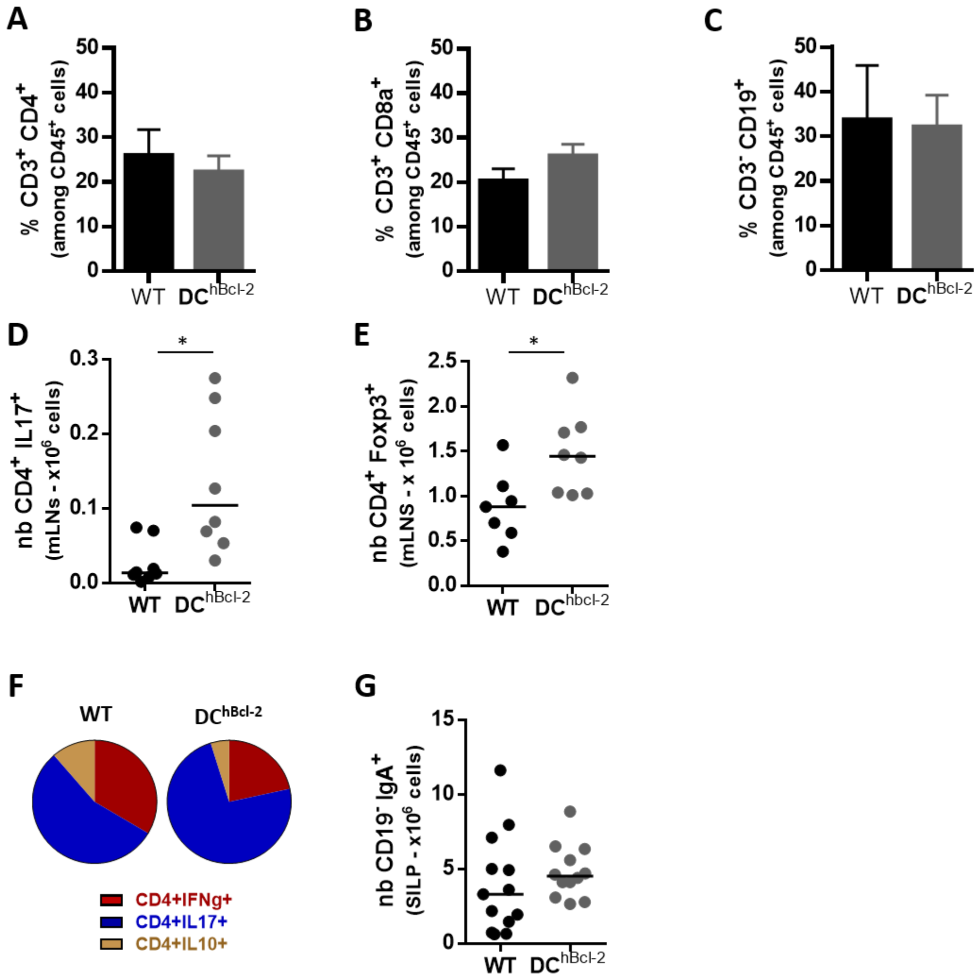


**figure supplement 4** All the data are representative of mice fed a HFD for 24 weeks. Proportions of $CD3^+$ $CD4^+$ T lymphocytes (A) or $CD3^+$ $CD8a^+$ T lymphocytes (B) or $CD3^-$ $CD19^+$ B lymphocytes after surface staining of cells in the mLNs (N=4 to 5 mice per group). Total numbers of IL-17-producing $CD4^+$ T (Th17) cells (D) or $CD4^+$ $Foxp3^+$ T (Treg) cells (E) after intracellular staining of cells in the mLNs. (F) Circle graphs representing the mean proportions of IFNg-producing, IL-17-producing, IL-10-producing $CD4^+$ T cells in the SILP after intracellular staining of cytokines. (G) Total numbers of $CD19^-$ $IgA^+$ plasmablasts after surface staining of cells in the SILP.

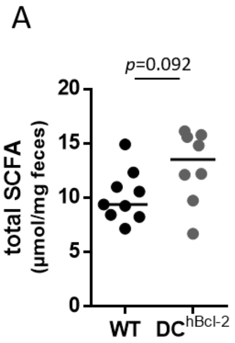


**figure supplement 5** Total short chain fatty acids (SCFA) concentration in the feces of mice after 24 weeks of HFD.

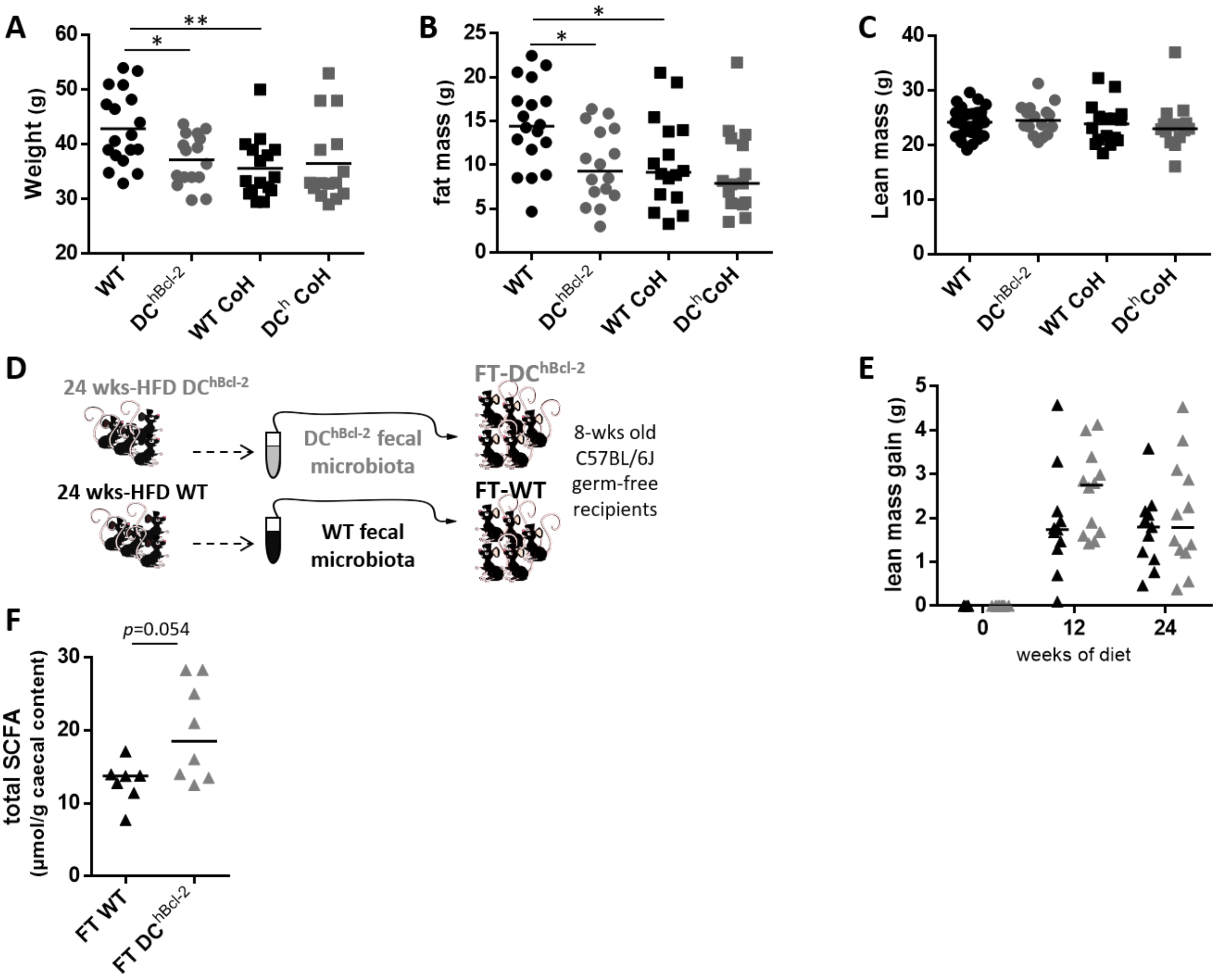


**figure supplement 6** Weight (A), fat mass (B) and lean mass (C) of single housed (dots) versus cohoused (squares) WT and DC$^{hBcl-2}$ mice after 24 weeks of HFD. (D) Fecal transplantation scheme from 24-weeks of HFD-fed donors into 8-weeks old germ-free recipients immediately submitted to HFD for 24 weeks. (E) Lean mass gain monitoring of recipients (FT)-mice at day 0, 12 weeks and 24 weeks after both fecal transplantation and starting the HFD. (F) Total short chain fatty acids (SCFA) concentration in the feces of FT-mice 24-weeks after both fecal transplantation and starting the HFD.
